# Impact of the yeast S0/uS2-cluster ribosomal protein rpS21/eS21 on rRNA folding and the architecture of small ribosomal subunit precursors

**DOI:** 10.1101/2023.01.18.524556

**Authors:** Gisela Pöll, Joachim Griesenbeck, Herbert Tschochner, Philipp Milkereit

## Abstract

RpS0/uS2, rpS2/uS5, and rpS21/eS21 form a cluster of ribosomal proteins (S0-cluster) at the head-body junction near the central pseudoknot of eukaryotic small ribosomal subunits (SSU). Previous work in yeast indicated that S0-cluster assembly is required for the stabilisation and maturation of SSU precursors at specific post-nucleolar stages.

Here, we analysed the role of S0-cluster formation for rRNA folding. Structures of SSU precursors isolated from yeast S0-cluster expression mutants or control strains were analysed by cryogenic electron microscopy. The obtained resolution was sufficient to detect individual 2’-O-methyl RNA modifications using an unbiased scoring approach.

The data show how S0-cluster formation enables the initial recruitment of the pre-rRNA processing factor Nob1 in yeast. Furthermore, they reveal hierarchical effects on the pre-rRNA folding pathway, including the final maturation of the central pseudoknot.

Based on these structural insights we discuss how formation of the S0-cluster determines at this early cytoplasmic assembly checkpoint if SSU precursors further mature or are degraded.

## Introduction

The small ribosomal subunit (SSU) of the yeast S. cerevisiae (hereafter referred to as yeast) consists of 33 ribosomal proteins (r-proteins) and the 18S ribosomal RNA (rRNA). R-proteins are subsequently named according to the standard yeast nomenclature, and additionally in the text according to the universal nomenclature [1].

In mature SSUs, each of the 34 components is folded in a highly defined way [2]. Moreover, an intricate network of protein-protein, protein-RNA, and tertiary RNA interactions results in a very stable overall tertiary structure while still allowing for some conformational flexibility during the different phases of mRNA translation. In the case of the 18S rRNA, 45 universally conserved helices are organized in four secondary structure domains termed 5’ domain, central domain, 3’ major domain and 3’ minor domain (see [3] for an overview). These are the main building blocks of dominant SSU tertiary structure features termed the ‘head’ (3’ major domain), and the ‘body’ (5’ domain, 3’ minor domain) with its ‘platform’ (central domain) which are already distinguishable at low-resolution. The body and the head are connected by the ‘neck’ helix h28 which arises from the body helix h2, also called central pseudoknot or CPK. In the CPK the terminal loop of the 5’ domain helix h1 base pairs with the rRNA region between the central domain helix h27 and the 3’ major domain ‘neck’ helix h28. The CPK is thus a ‘central’ RNA-based connecting point in the SSU which restraints the spatial distance between the 5’, the central, and the 3’ major domain.

Formation of the SSU in yeast cells, as in other organisms, is a highly energy-demanding process. It involves besides the folding and assembly of the structural components other aspects such as the synthesis and intracellular transport of these components, and the modification of specific rRNA residues (reviewed in [4–6]). Moreover, the 18S rRNA is synthesized in the nucleolus as part of a large precursor RNA (pre-rRNA). Here, it is flanked on the 5’ site by the 5’ external transcribed spacer (5’ ETS) and on the 3’ site by the internal transcribed spacer 1 (ITS1). Consequently, these have to be removed during SSU maturation (see [7,8] for an overview). Removal of the 5’ ETS still occurs in the nucleus and thus results in a nuclear 20S pre-rRNA containing SSU precursor population. After nuclear export to the cytoplasm, the ITS1 is cleaved off at one point by the PIN-domain endonuclease Nob1 [9–12]. Besides the nucleases and RNA modifying enzymes involved in pre-rRNA processing and modification, many other factors were identified that transiently interact at specific stages with the maturing SSU precursors. Cryogenic electron microscopy analyses (cryo-EM) of particles affinity purified from cells via tagged versions of such factors provided several structural snapshots which possibly represent folding intermediates of the nascent maturing SSU (see [13] for an overview).

Based on combined structural data obtained from yeast and human SSU precursor particles, a possible timeline of events can be deduced by ordering snapshots according to the stepwise appearance of features observed in mature SSUs. Accordingly, at an early nucleolar stage, the 5’ ETS serves as a platform for the recruitment of numerous factors (reviewed in [6,14,15]). These in turn interact with SSU components and, among other aspects, keep in the resulting large particles the nascent SSU secondary structure domains spatially separated from each other. Among them is the U3 small nucleolar RNA (snoRNA) which directly prevents the formation of the CPK by base pairing with the respective SSU rRNA regions. At later, presumably nucleoplasmic stages, the 5’ ETS, the U3 snoRNA and most of the early associating factors have been released [16–21].

Instead, a new group of factors associates with the SSU precursor. Combined biochemical and structural data support that this group includes the endonuclease Nob1, Tsr1, Rio2, Bud23, Slx9, and Rrp12 together with Pno1, Dim1, and Enp1 which are retained from previous states [18,22]. Different from their human counterparts [23–25], yeast Nob1, Slx9, and Rrp12 were barely detected at any structural snapshot [26–30]. Still, Rrp12 is thought to facilitate in yeast and human at this maturation stage the passage through the nuclear pore by direct interaction with pore components [31]. In addition, Slx9 and Rio2 were suggested to further support the nuclear export of these SSU precursors by recruiting the exportin Crm1 [32]. Not only the factor composition, also the r-protein assembly state and fold of SSU precursors changes after the release of the 5’ETS, early factors and of the U3 snoRNA. The CPK was observed in early cytoplasmic SSUs in a mature conformation concomitant with a mature-like head-body arrangement [24,26–28]. The neck helix h28 is still slightly distorted at this point, which results in an atypical head to body tilt. In addition, a cluster of the three r-proteins rpS0/uS2, rpS2/uS5, and rps21/eS21 (subsequently termed S0-cluster) is now detected near the newly formed CPK and, in human particles, near Nob1. S0-cluster formation might precede Nob1 integration, as suggested by a structural state unexpectedly observed in human particles associated with a dominant negative mutant form of the late acting factor human RIOK1 (named Rio1 in yeast) [22]. The wild-type form of RIOK1 is only detected in late cytoplasmic SSU precursors after the release of most of the previous factors. At this point, also a cluster of r-proteins at the helix h33 and an adjacent protein cluster around uS3 [29,33] are newly formed in a mature-like conformation at the ‘beak’ of the SSU head [23,24].RIOK1/Rio1 is thought to facilitate in the platform of these late cytoplasmic SSU precursors together with rpS26/eS26 the release of Pno1 near the 18S rRNA 3’ end. This then enables the Nob1 endonuclease active site to access the 18S rRNA 3’ end and to split off the ITS1 [23,25,34– 36].

Altogether, the arrangement of the observed structural snapshots in such a timeline suggests that SSU assembly and folding proceeds in principle in a series of consecutive events which is accompanied by recruitment and release of specific factors. Still, alternative, and parallel assembly pathways might have been missed due to the possibly short-lived nature of respective intermediates.

Importantly, it cannot be unambiguously distinguished based on the deduced timeline of structural snapshots if a structural feature is observed before another one merely because of different kinetics to establish them, or due to a true hierarchical relationship. Such hierarchical interrelationships between events potentially have important functional consequences. They can streamline the maturation process and introduce quality control mechanisms by favouring the incorporation of factors and r-proteins into SSU-precursors with defined maturation state. On the other hand, further maturation of particles with specifically misassembled or misfolded regions may be blocked, thus exposing them to general degradation pathways over time. The first aspect can be especially favourable if only a limited free pool of these components is available, of which the r-proteins have to be produced in growing cells in considerable amounts. Moreover, there are even indications that large excess of free r-proteins can lead to their aggregation and toxic effects on cellular protein folding homeostasis [37–39].

To gain more insights into functional interrelationships between r-protein assembly and pre-rRNA folding events, we analysed here the impact of the assembly of the S0-cluster of r-proteins on the pre-rRNA fold and the general architecture of late nucleoplasmic to early cytoplasmic yeast SSU precursors. Previous biochemical studies indicated that S0-cluster formation is required for the production and accumulation of yeast SSUs [33,40–45]. Effects were observed on the dynamics of their nuclear export, the cytoplasmic release of the ITS1, the recruitment of factors such as Nob1, and the release of other factors such as Slx9, Rrp12, and Bud23. To visualize the impact on the nascent SSU folding state we compared here by cryo-EM analyses the structure of the respective particle populations purified from a control or a mutant yeast strain in which formation of the S0-cluster was inhibited.

## Results

### Purification of yeast Enp1-associated SSU precursors after expression shutdown of the S0-cluster r-protein rpS21/eS21

To analyse the role of the S0-cluster protein rpS21/eS21 for the folding of SSU precursors we made use of a conditional lethal mutant yeast strain in which rpS21/eS21 is expressed under the control of the galactose inducible GAL1/10 promoter [33,44]. The genome of this strain and the one of a control strain in which rpS21/eS21 is expressed from its endogenous promoter was further modified to encode for the essential SSU biogenesis factor Enp1 in fusion with the tandem affinity purification tag (TAP-tag) (see Materials and Methods). Enp1 initially associates with nucleolar SSU precursors already in the nucleolus and dissociates from them in the cytoplasm [46]. It is detected in cryo-EM-based structural snapshots of yeast SSU precursors before the appearance of S0-cluster r-proteins and leaves at a stage at which they are found at near-mature positions.

Both strains were cultivated for four hours in a glucose-containing medium to shut down the expression of rpS21/eS21 in the mutant strain. Affinity purification of TAP-tagged Enp1 and associated particles were then performed with cellular extracts of the two strains. Northern blotting experiments confirmed that 20S pre-rRNA containing SSU precursors co-enriched with TAP-tagged Enp1 from control and rps21 mutant extracts (S1 Appendix A and B, lanes 5 and 6). The observed increased ratios of 20S pre-rRNA to 18S rRNA and of 25S rRNA to 18S rRNA after expression shutdown of rpS21/eS21 were also consistent with a role of S0-cluster r-proteins for efficient processing of 20S pre-rRNA and accumulation of SSUs (S1 Appendix A-C) [41–45].

### Detection of 2’-O-methyl adenosine residues in a cryo-EM derived density map of Enp1-associated SSU precursors by an unbiased scoring approach

The two affinity-purified samples were subsequently analysed by single particle cryo-EM to deduce major structural features of the contained SSU precursors (see Materials and Methods). Enp1-TAP associated SSU precursors from the control strain were predominantly found in a conformational state subsequently termed Enp1TAP_A (see S2 Appendix) for which a corresponding pseudo-atomic model was built (see Materials and Methods). The global Enp1TAP_A map resolution was estimated to surpass 3 Å (see S2 Appendix and Materials and Methods). We thought to further examine the map and model quality by testing in an unbiased way for known 2’-O-methyl modifications of adenosine residues in the 18S pre-rRNA. For this, all adenosines in the model Enp1TAP_A 18S rRNA were converted into 2’-O-methyl adenosines and a global molecular dynamics simulation of the modified model with the given Enp1TAP_A map was performed. The map value for the newly introduced methyl groups at each adenosine position was then determined and used as a first scoring criterion (Sc_MapVal). To probe for an expected falloff in map values further away from the predicted methyl group position, the length of the given bonds between the 2’-O atom and the carbon atom of the methyl group was extended to 2.4 Å. The ratio of map values at the originally predicted and the more distant positions was calculated for each methyl group as a second scoring criterion (Sc_FallOff). When the >400 given adenosines in model Enp1TAP_A were sorted according to the Sc_MapVal criterion, all of the 8 known 18S rRNA 2’-O-methyl adenosines ranked among the top 15 (see S3 Appendix, A). Further inspection showed that these known 2’-O-methyl adenosines all had Sc_FallOff values higher than 1.5 while 5 out of the 7 other top 15 ranked adenosines had clearly lower Sc_FallOff values (see S3 Appendix, A). Manual inspection indicated that in these cases, some unattributed density of for example ions was close to the hypothetical methyl groups after the molecular dynamic’s simulation.

From this unbiased scoring result, we conclude that individual methyl groups start to be resolved in significant parts of the Enp1TAP_A map. When combined with subsequent manual inspection, similar approaches to the one utilized here can potentially be applied to screen for other nucleotide or amino acid residue modifications in an unbiased and automatable way. As indicated by the described scoring experiments, the modifications of interest can be as small as methyl groups if the provided map and model are of sufficient quality.

### Overall architecture of the Enp1TAP_A SSU precursor

Key aspects of the overall architecture of the Enp1TAP_A SSU precursor model resembled the ones previously found for yeast and human late nucleoplasmic and early cytoplasmic SSU precursors [24,26–28]. Factors specifically binding at early nucleolar stages of SSU maturation as well as the U3 snoRNA and associated proteins were not detected. Conversely, base pairing in the central pseudoknot as seen in mature SSUs could well be traced in Enp1TAP_A. In addition, all r-proteins (except rpS26/eS26 in the platform at the 18S rRNA 3’ end) binding to the SSU body, including the S0-cluster r-proteins, as well as factors previously observed in yeast early cytoplasmic SSU precursors were found at their expected positions (Rio2, Enp1, Pno1, Tsr1) (see Fig 1 A for an overview). Several parts of the 3’ major domain which are thought to mature at later cytoplasmic stages were still found in an ill-defined state. These included helices h31, h34, h40, and an incomplete and distorted ‘neck’ helix h28, the latter resulting in an immature head-to-body orientation. Similarly, a cluster of proteins at the beak helix 33 (r-proteins rpS10/eS10, rpS12/eS12 and rpS31/eS31) together with the adjacent protein cluster formed by rpS3/uS3, rpS20/uS10, and rpS29/uS14 could not yet be detected at the SSU-head.

**Fig 1.**
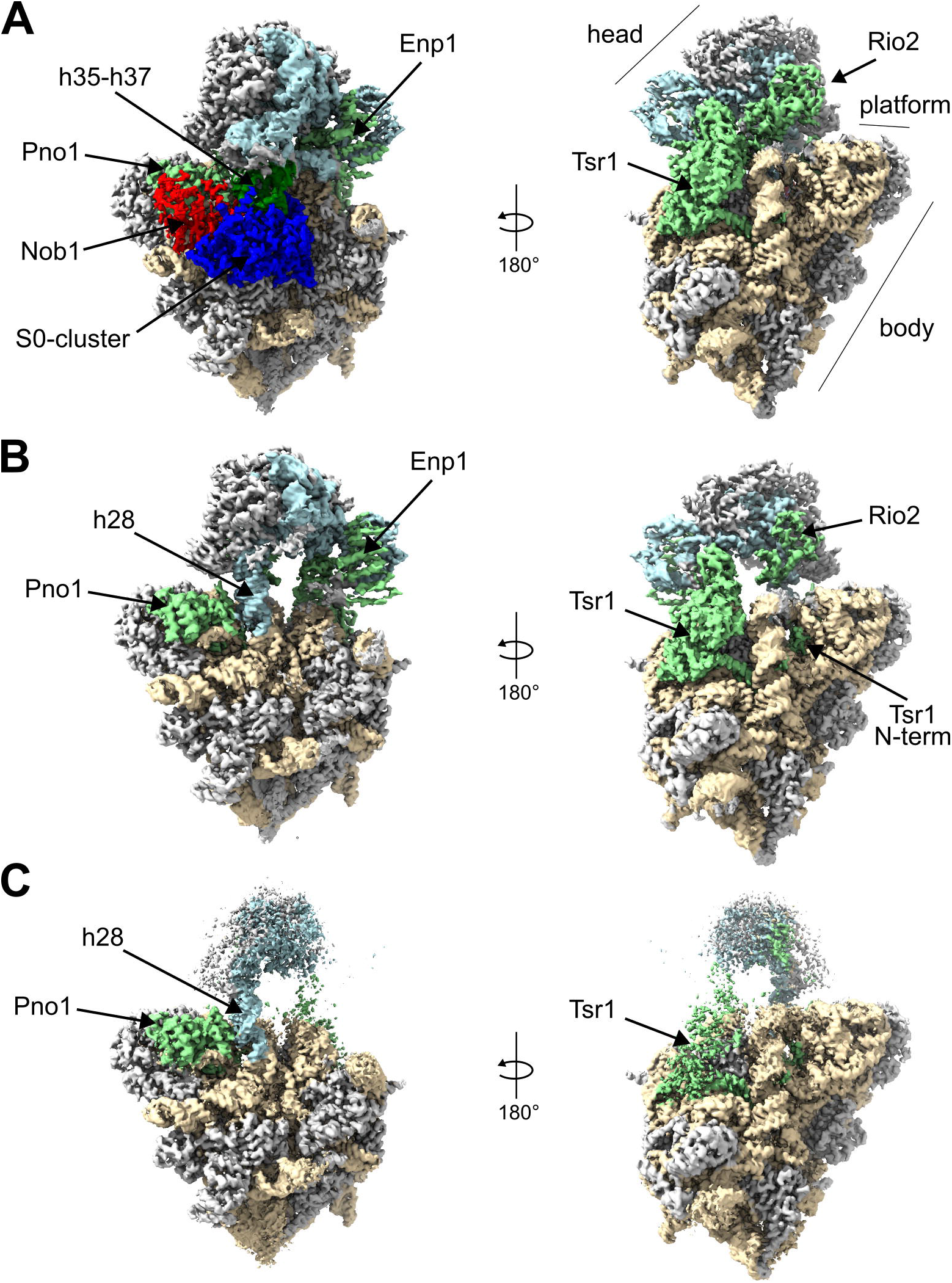
Cryo-EM derived maps of SSU precursor populations Enp1TAP_A (A), Enp1TAP-S21_A (B), and Slx9 TAP-S21_A (C). Helices h35-h37 are shown in dark green, other rRNA of the head in light blue, rRNA of the body in light brown, the S0-cluster in dark blue, Nob1 in red, and all other factors in light green, and r-proteins in gray.

Besides, at the given map resolution more detailed insights were revealed in regard to the structure of the analysed yeast early cytoplasmic SSU precursor population. Among them were the binding mode of yeast endonuclease Nob1 to these SSU precursors and the pre-rRNA path downstream of the cleavage site at the 18S rRNA 3’ end (see further below).

The obtained map also provided evidence for an unexpected conformation of the terminal loop of helix h45. In mature ribosomes, the terminal loop of h45 is arranged in a tetraloop (U1779-A1782) conformation with pi-stacking of bases G1780, A1781, and A1782 (see Fig 2B). The N6 atoms of A1781 and A1782 are each dimethylated in mature ribosomes and it is thought that the dimethylation is finalized in yeast by the universally conserved enzyme Dim1 on cytoplasmic SSU precursors [47–49]. Remarkably, the observed density for the H45 loop supports its arrangement at this stage as hexaloop. The bases of nucleotides A1782 and C1783 are aligned in a possible stacking configuration perpendicular to the two adjacent stacked bases of nucleotides G1780 and A1781 (see Fig 2A). The local resolution was not sufficient to unambiguously determine if the N6 atoms of residues A1781 and A1782 were already dimethylated by Dim1. Still, previous biochemical analyses of SSU precursors purified via Enp1-Tap indicated an incomplete methylation state of nucleotide A1781 at this stage [49]. Reminiscent of this, only partial methylation at the adenosine corresponding to A1781 was recently observed in mature ribosomes of some archaeal species where the G-C base pair following the tetraloop is converted into a weaker G-U pair [50].

**Fig 2.**
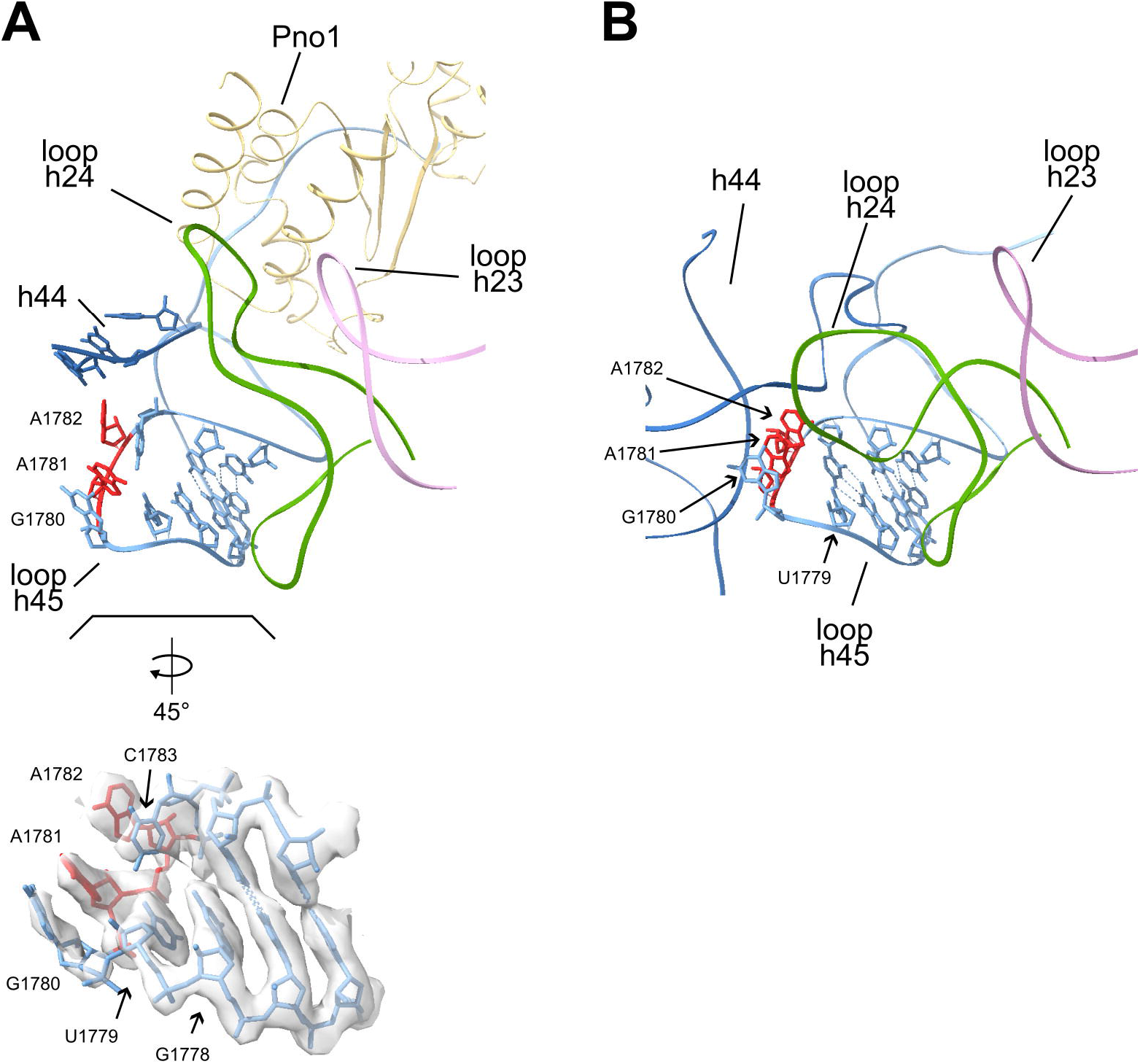
Structure model of helix h45 and its context in the Enp1TAP_A SSU precursor population (A) and in mature ribosomes (B). The backbone of helix h45 is shown in light blue, and the ones of h44, h23, and h24 are in dark blue, pink, and green, respectively. For nucleotides in the terminal loop of helix h45 atomic models are shown, with the substrate nucleotides of Dim1 (A1781 and A1782) in red. The lower insert in (A) shows the terminal loop of helix h45 further enlarged, rotated by 45 degrees, and overlayed with the cryo-EM derived map of Enp1TAP_A SSU precursors. Pdb entry 4v88 was used for the visualization of mature ribosomes.

While the detected premature orientation of helices h24 and h44 keeps the terminal loop of helix h45 still accessible in Enp1TAP_A particles (compare helices h24 and h44 in Figs 2A and 2B) we did not observe any clear density corresponding to Dim1 there or elsewhere. Dim1 was previously found in proteomic studies as a prominent component of Enp1-TAP associated yeast SSU precursors [46]. It might therefore bind at this stage to still flexible parts of the SSU precursor, obscuring its visualization by cryo-EM analyses. In support of this, previous UV-crosslinking analyses identified a major rRNA binding site of Dim1 in the 3’ part of helix h44 which could not be resolved in Enp1TAP_A particles [51].

### Structure of the yeast endonuclease Nob1 and interacting internal transcribed spacer rRNA in Enp1TAP_A particles

During processing of the yeast Enp1-TAP particle dataset, we observed at the SSU platform between Pno1 and the S0-cluster r-proteins a previously unassigned density at the position where Nob1 is located in human SSU precursors [24]. Similar to its human counterpart, yeast Nob1 consists of an N-terminal PIN-domain [10] with an insertion domain, followed by a zinc ribbon, and a C-terminal domain [11,52]. A model generated by the AlphaFold algorithm [53,54] of the yeast Nob1 PIN-domain and the zinc ribbon could be unambiguously fitted into parts of the unassigned density. Otherwise, the rest of the unassigned density could be attributed to parts of the Nob1 insertion domain and the C-terminal domain which partially deviated regarding their detailed fold or orientation from the AlphaFold predictions. Yeast Nob1 main contacts with the SSU precursor were found to be established in Enp1TAP_A particles by interactions of the insertion domain with Pno1, of the zinc finger with rpS0/uS2, and of the C-terminal domain with rpS0/uS2 and rRNA helices h26, h36 and h37 in the SSU platform (see Figs 3A and 3C). These regions of Nob1 were in general better resolved than its PIN-domain which was detected here on top of the zinc ribbon and the insertion domain. The observed interactions of yeast Nob1 with helices h26 and h37 were consistent with previous UV-crosslinking analyses [35]. The active center in the PIN-domain was observed far from its substrate, the region around the 18S pre-rRNA 3’ end. This “inactive” constellation of Nob1 resembles the one observed in human early cytoplasmic SSU precursors [24]. Still, the orientation of the PIN domain towards the zinc ribbon differed between the yeast and human early cytoplasmic SSU precursor models by a rotation of about 35°.

**Fig 3.**
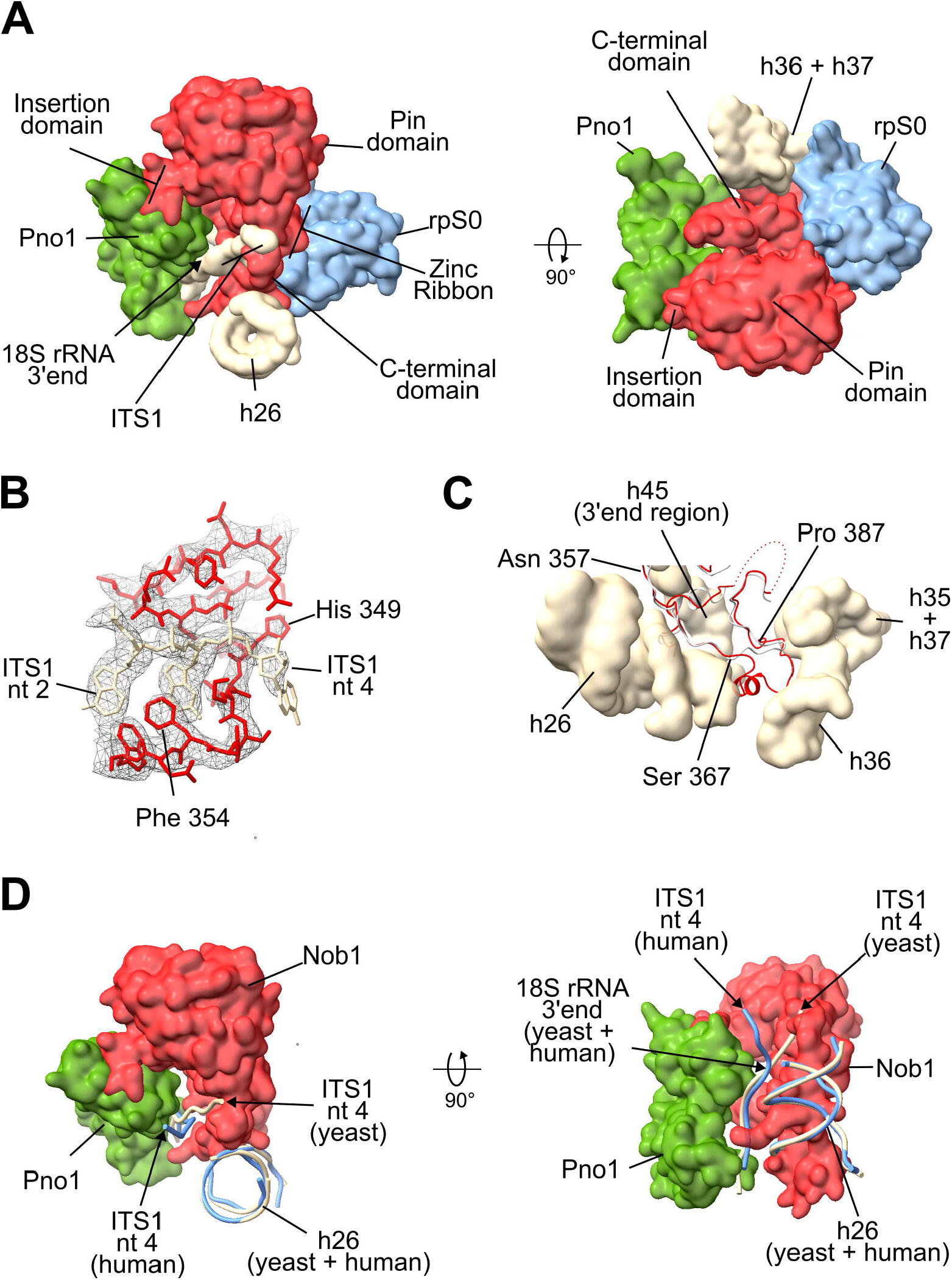
Structure model of yeast Nob1 in early cytoplasmic SSU precursors. (A) shows an overview of Nob1 together with its interaction partners, (B) the region around ITS1 nucleotides 2-4 and (C) the yeast and human (gray) C-terminal domains of Nob1 in the context of rRNA helices h26, and h35-h37. In (D) the path of human (light blue) and yeast pre-rRNA around the 18S rRNA 3’ end is compared. In (A) - (D) yeast Nob1 is shown in red, Pno1 in green, rpS0/uS2 in blue and pre-rRNA in wheat. In (A), (C) and (D) the Enp1TAP_A model is visualized by surface representations, except for the backbone representation of Nob1 in (C) and pre-rRNA in (D). In (B) the full atomic model is shown together with the cryo-EM derived map of Enp1TAP_A SSU precursors. Pdb file 6g18 was used for the visualization of human Nob1 in early cytoplasmic SSU precursors ().

The obtained density map for the Enp1TAP_A state also allowed to trace the rRNA precursor backbone for 4 nucleotides downstream of the mature 18S rRNA 3’ end into the ITS1. Clear densities for the adenine base at ITS1 position +2 and the guanine base at position +3 were observed while the orientation of bases at the 18S rRNA 3’ end and at ITS1 positions +1 and + 4 was less defined. Accordingly, yeast ITS1 pre-rRNA passes along the beta-sheets of the Nob1 zinc ribbon domain and crosses at residues +3 and +4 the Nob1 peptide backbone between histidine 349 and phenylalanine 354 (Fig 3B). That is the region, where the Nob1 C-terminal domain emerges from the zinc ribbon domain (Fig 3A). This path is clearly different from the one observed for the first four ITS1 nucleotides in human early cytoplasmic SSU precursors, which are more oriented towards Nob1’s interaction partner Pno1 (Fig 3D). Still, in both cases, the Nob1-Pno1 interface can potentially protect at this maturation stage the single-stranded SSU pre-rRNA around the 18S rRNA 3’ end and 4 nucleotides beyond from degradation by endo- and 3’ - 5’ exonucleases.

### The initial positioning of Nob1 and rpS17/eS17 in early cytoplasmic SSU precursors depends on S0-cluster assembly

We next analyzed the cryo-EM dataset of the Enp1-associated SSU precursors which were purified from cells in which expression of the S0-cluster r-protein rpS21/eS21 was shut down. Two major populations with distinct folding states could be distinguished after particle sorting and three-dimensional structure reconstruction (S4 Appendix). They are termed in the following Enp1TAP-S21_A, to which about 132.000 particles contributed, and Enp1TAP-S21_B with about 69.000 particles. The Enp1TAP-S21_B map was highly similar to the Enp1TAP_A map from control cells and showed clear density of rpS21/eS21 and the other S0-cluster proteins. We consider that the corresponding particles were purified from cells with ongoing residual expression of rpS21/eS21.By contrast, in the Enp1TAP-S21_A map, no density corresponding to rpS21/eS21 or the other S0-cluster r-proteins could be detected, consistent with previous evidence for their cooperative stable integration into the SSU [33]. Otherwise, the overall architecture of the derived Enp1TAP-S21_A model was similar to the Enp1TAP_A control model (Fig 1B, compare with Enp1TAP_A in Fig 1A) with several exceptions detailed below. The estimated global resolution of the Enp1TAP-S21_A map was estimated to be close to 3 Å (S4 Appendix and Materials and Methods), and again, unbiased scoring of 2’-O-methyl adenosine residues indicated that individual methyl groups started to be resolved in significant parts of the map (see S3 Appendix B, 7 out of 8 known 2’-O-methyl adenosines in the top 20 of >400 adenosine residues).

No indication was observed for the presence of the two r-protein cluster interacting proteins rpS17/eS17 and Nob1 at their expected positions. These observations matched with previous proteomic analyses of rpS21/eS21 depleted SSU precursors pointing to a role of S0-cluster assembly for stable recruitment of rpS17/eS17 and Nob1 [33]. The large interaction interface of the S0-cluster with these proteins (see above and interaction analyses for the Enp1TAP_A model in S5 Appendix) is likely directly responsible for the establishment of the observed binding hierarchy.

The average map values of Rio2 were reduced by about 30% in the Enp1TAP-S21_A map when compared to the Enp1TAP_A map from the control strain (see Materials and Methods). Average map values of Pno1 and the bait protein Enp1TAP remained on similar levels (both <5% changes). The Rio2 N-terminal region was hardly visible, consistent with a supportive role of S0-cluster formation for the correct incorporation of Rio2 into SSU precursors [33]. For the factor Tsr1, a part of its N-terminal domain could be modelled into densities between SSU rRNA helices h19, h27, and h45. Corresponding densities were only scarcely visible in the Enp1TAP_A map. This newly mapped N-terminal portion of yeast Tsr1 partly overlaps with the recently observed position of the human Tsr1 N-terminus, but the described extension towards the immature helix h28 was not detected in the Enp1TAP-S21_A map [22].

### Association of the export factor Slx9 with a sub-population of rpS21/eS21 depleted SSU precursors with flexible head architecture

Previous analyses indicated that the release of the two factors Rrp12 and Slx9 from early cytoplasmic SSU precursors is delayed when S0-cluster assembly is inhibited [33,40]. Both factors were previously implicated in the nuclear export of yeast SSU precursor particles [31,32]. No clear density could be attributed to these factors in the Enp1TAP-S21_A map. The same was true when SSU precursor particles were purified via Slx9-TAP from cells in which expression of rpS21/eS21 was inhibited (S1 Appendix A and B, lanes 4 and 8). Here, cryo-EM analyses with subsequent particle sorting and reconstruction of three-dimensional maps (S6 Appendix) indicated that for the main population of SSU-precursors (termed Slx9TAP-S21_A) the global architecture of the SSU body resembled the one found for the Enp1TAP-S21_A population (Fig 1C, compare with Fig 1B), including the lack of the S0-cluster, Nob1, and rpS17/eS17. Still, the density for the SSU head was clearly less defined in the Slx9TAP-S21_A map. Accordingly, yeast Slx9 binds to a rather small subpopulation of rpS21/eS21-depleted SSU precursors (compare signals in lanes 6 and 8 in S1 Appendix, A) with comparably large flexibility in the SSU head orientation or architecture.

### Impact of S0-cluster assembly on the stable orientation of rRNA helices h35 - h37 and on final maturation of the CPK

The general pre-rRNA fold observed in the Enp1 TAP-S21_A map largely resembled the one of the Enp1TAP_A map with a few exceptions. The three RNA helices h35 - h37 were not detectable any more in Enp1TAP-S21_A (Fig 1B, compare with Enp1TAP_A in Fig 1A). These were well resolved in the Enp1TAP_A map, flanked by the CPK and the neck-helix h28 on one side, and by the S0-cluster and the Nob1 C-terminal domain on the other side. Apparently, the establishment of a defined orientation of these helices depends on S0-cluster assembly. Direct interactions of helices h35 - h37 with S0-cluster r-proteins and with the downstream recruited Nob1 (S7 Appendix, see predicted interactions of helices h35 - h37 in Enp1TAP_A, see also Nob1 interactions with helices h35 - h37 in Figs 3A and 3C) possibly play a direct role in this process.

More detailed inspection showed that there were still fewer base pairs established in the body-proximal part of the neck-helix h28 of Enp1TAP-S21_A particles compared to Enp1TAP_A particles or mature ribosomes (see secondary structure diagrams in Figs 4B-4D for an overview). Further analyses of the map indicated that the helix h28 nucleotide U1145 was incorporated into an immature form of the adjacent CPK helix h2, which we subsequently term CPK-im (Figs 4A and 4D). U1145 forms in mature ribosomes with nucleotide A1633 the terminal base pair of the body-proximal part of helix h28. In Enp1TAP-S21_A, nucleotide U1145 replaced U1144 in forming the first wobble base pair in helix h2 with G10, causing thereby a register shift, and allowing nucleotide U1144 to base pair with nucleotide A11. The base of nucleotide U9 was observed reoriented in CPK-im towards nucleotide A1143, thus establishing a new base pair between U9 and A1143 (Figs 4A and 4D). The series of first base interactions of the mature CPK, U1144:G10, A1143:A11, and A1142:U12 was thereby converted in CPK-im into U1145:G10, U1144:A11, A1143:U9, and A1142:U12. Formally, the latter arrangement should provide higher paring energy in helix h2 than the first one. All following base pairs in CPK-im resembled the one of the mature CPK helix h2.

**Fig 4.**
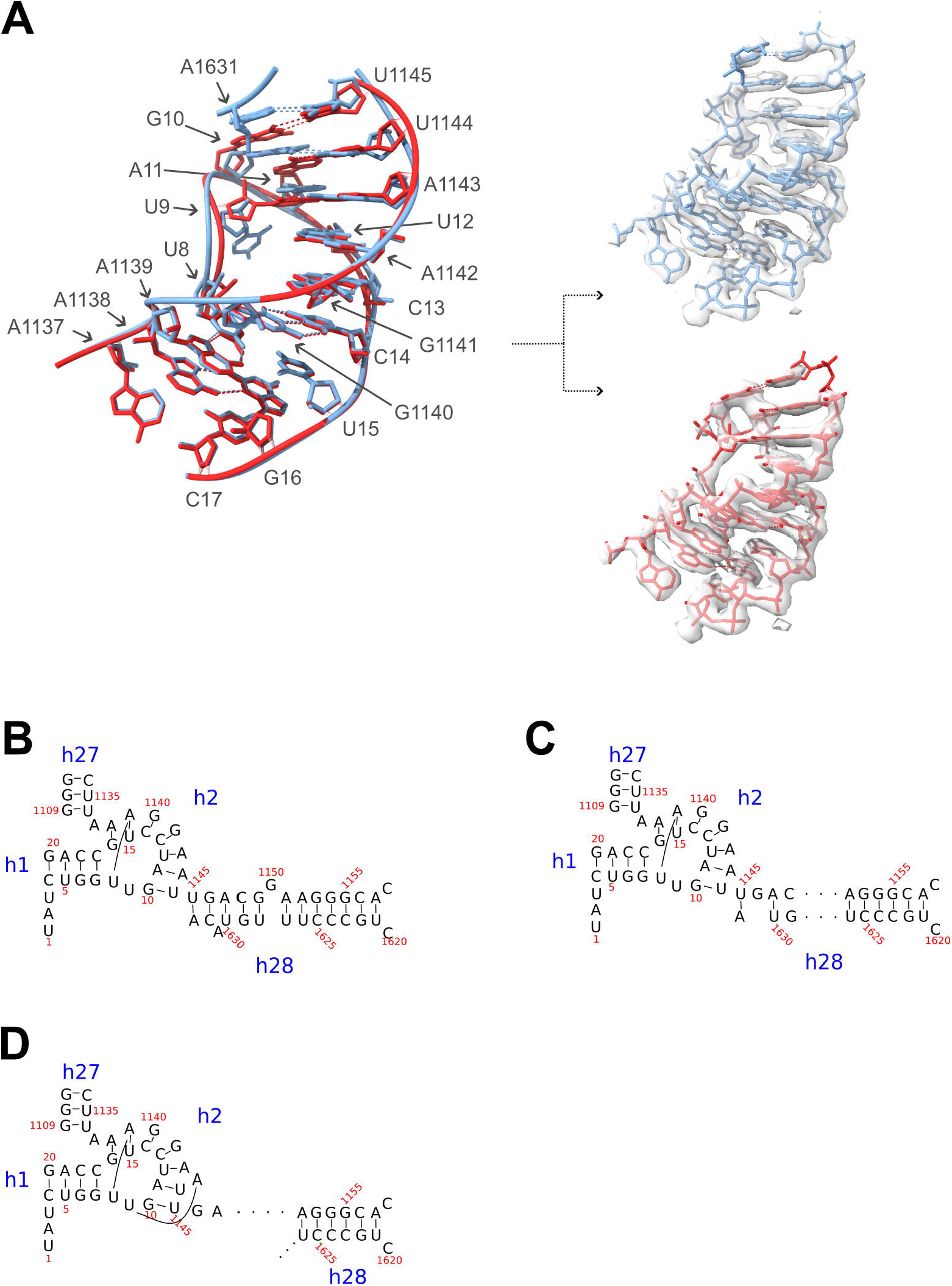
Structure model of the CPK-im in Enp1TAP-S21_A compared to the mature CPK in Enp1TAP_A particles. In (A) on the left, the structure models of CPK-im (red, Enp1TAP-S21_A particles) and the mature CPK (light blue, Enp1TAP_A particles) are superimposed. On the right, they are shown individually with the respective overlaid cryo-EM derived maps Enp1TAP-S21_A and Enp1TAP_A. In (B) - (D) secondary structure models are shown of the CPK in yeast mature ribosomes (B), in Enp1TAP_A particles (C), and in Enp1TAP-S21_A particles (D).

We were interested to find out if the formation of CPK-im depends on the specific situation induced by expression shut down of rpS21/eS21 or if this state can also be found in cells with normal expression levels of rpS21/eS21. In the latter case, it possibly represents a metastable intermediate during yeast CPK formation whose further maturation depends on the progressive assembly of the S0-cluster. To test this, further particle sorting was performed on the Enp1TAP dataset from control cells, focussing on the region containing the S0-cluster. Indeed, a proportion representing about 9% of the total identified SSU precursor particles fell into a class of particles lacking the S0-cluster, subsequently termed Enp1TAP_B (S2 Appendix). The overall SSU precursor architecture deduced from the Enp1TAP_B map highly resembled the one observed for Enp1TAP-S21_A particles depleted of rpS21/eS21 (compare Fig 1B with Fig 5A). Again, densities for the S0-cluster, rpS17/eS17, Nob1, and helices h35-37 could not be detected, while other global architectural SSU features found in the Enp1TAP_A particle population were preserved. Still, a more detailed inspection indicated that helix h2 was in the CPK-im state with U1145:G10, U1144:A11, and A1143:U9 base pairing (Figs 5B and 5C). Thus, CPK-im could be detected in a subpopulation of particles isolated from cells with endogenous expression levels and ongoing assembly of the S0-cluster r-proteins. Altogether, we take these observations as evidence for S0-cluster assembly being required for and preceding the final maturation of the CPK.

**Fig 5.**
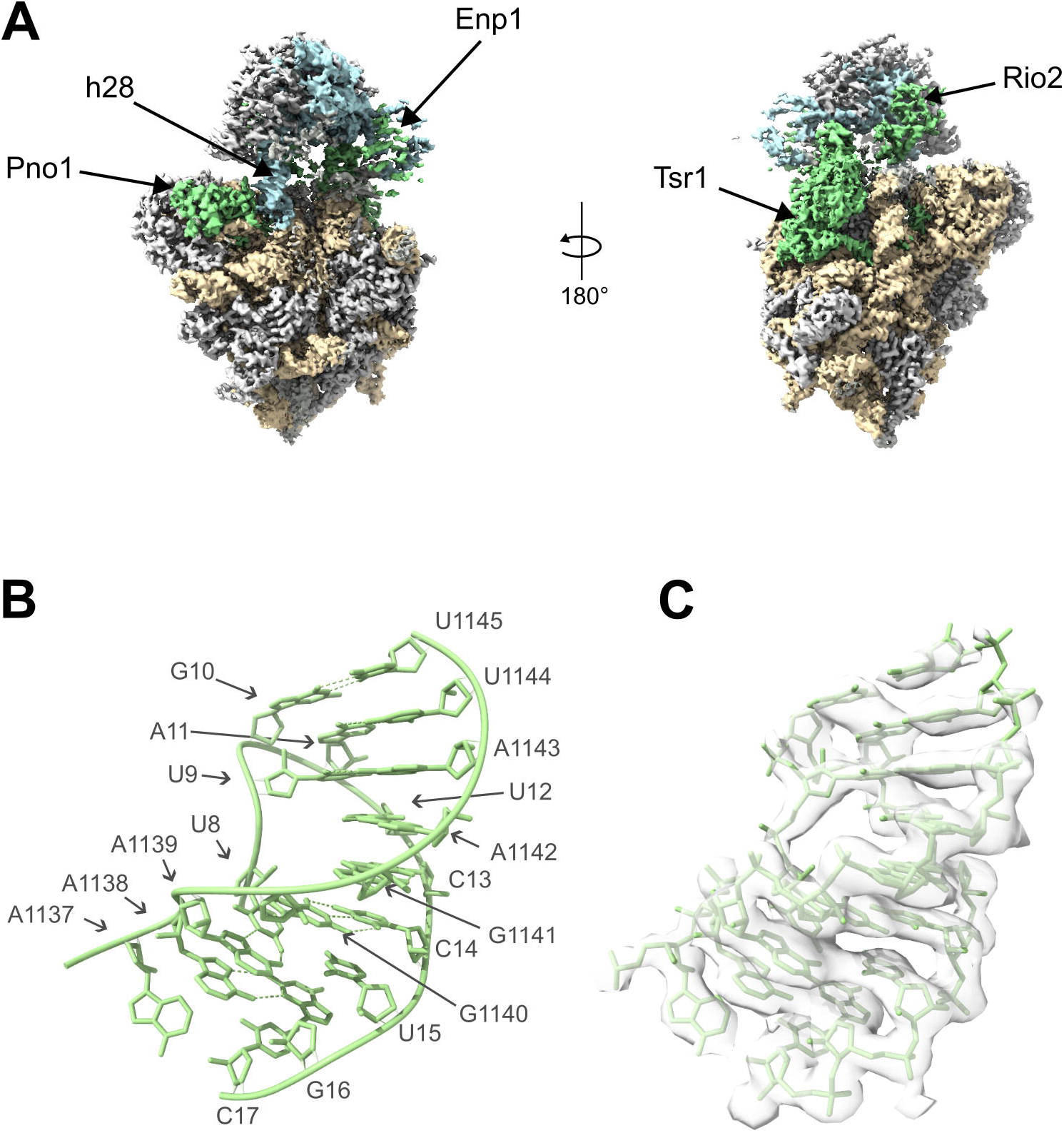
Incomplete S0-cluster assembly and CPK maturation in the Enp1TAP_B sub-population of SSU precursors. In (A) the cryo-EM derived map of the Enp1TAP_B particles is shown, in (B) the structure model of the CPK in these particles, and in (C) the structure model overlaid with the cryo-EM derived map of Enp1TAP_B particles. Coloring in (A) is done as described in the legend of Fig 1.

## Discussion

The data obtained give detailed insights into the hierarchical interrelationship between S0-cluster assembly and folding of nascent yeast SSUs. The binding interface between the S0-cluster r-protein rpS0/uS2 with yeast Nob1 was observed in near-atomic detail in Enp1TAP_A particles from control cells with the ongoing assembly of the S0-cluster. In the particles derived from the rpS21/eS21 expression mutant, not only rpS21/eS21 but also rpS0/uS2 and Nob1 were not detected. This suggests that formation of this binding interface during cooperative assembly of the S0-cluster is required for initial Nob1 incorporation. It is still possible that Nob1 can already interact independently of the S0-cluster via Pno1 or platform rRNA helices with SSU precursors, but this interaction seems not to be sufficient for its stable initial integration. These observations regarding mechanisms of Nob1 recruitment also further rationalized why the Nob1-dependent cut at the 18S rRNA 3’ end is affected in expression mutants of any of the S0 cluster r-proteins [42–45].

In addition, the yeast Nob1 structure in Enp1TAP_A particles showed that, like in human SSU precursors, Nob1 together with Pno1 can potentially shield at the early cytoplasmic stage the 18S pre-rRNA until base 4 of the ITS1 from endo- and exonucleolytic degradation. Indeed, indications for the degradation of accumulating 20S pre-rRNA were observed after the in vivo depletion of Nob1 [11]. Interestingly, Inhibition of general exosomal 3’-5’ degradation by inactivation of the Mtr4 helicase in yeast revealed that > 25% of 20S pre-RNA can be tagged in this situation by polyadenylation for the degradation via the exosome [55]. That indicates that the exosomal turnover of these SSU precursors can be substantial, especially upon failure of S0-cluster dependent shielding mechanisms. Besides, independent of Nob1 integration, the lack of S0-cluster assembly renders a large region around the CPK and the neck helix, including helices h35 - h37, more accessible to nucleases. The combined S0-cluster dependent shielding effects provide thus possible explanations for the decreased accumulation of SSUs in S0-cluster expression mutants [41–45].

We could not visualize the mode of binding of the three factors Slx9, Rrp12 and Bud23 whose release was previously observed to be affected by lack of S0-cluster assembly [33,40]. In Slx9-associated particles Slx9TAP-S21_A the SSU head was barely detectable. We consider that this specific, apparently small sub-population of S0-cluster depleted Slx9 associated SSU particles additionally contains Rrp12 and Bud23. The human counterparts of all three proteins interact with the SSU head [22], and their presence could thereby be obscured in the Slx9TAP-S21_A particle population due to the ill-defined head domain orientation or architecture.

The analysed structures furthermore provided evidence for S0-cluster assembly being required for the final maturation of the yeast CPK from an immature state CPK-im into the universally conserved state found in mature ribosomes. In CPK-im the last body-proximal residue of the incompletely formed neck helix h28 is incorporated into the CPK helix h2 and thus induces a frameshift in helix h2. In yeast, this frameshift is reversed at the third position by the formation of a new base pair with nucleotide U9 which is unpaired in the canonical CPK. Reinspection of recent cryo-EM derived maps and models of human late nuclear to early cytoplasmic SSU precursors indicates that the human CPK also exists in an immature state with the very same frameshift [22]. In this case, the frameshift is reversed by an unpaired nucleotide at position 3, which is different from the new base pair formed at this site in yeast with nucleotide U9. This human CPK-im state can be observed after the release of the U3 snoRNA (Rrp12-A states in [22]), right before the establishment of a stable orientation of the S0-cluster and the downstream neighboring elements in later states (Rrp12-B1 state in [22]). It is therefore possible that the S0-cluster dependent final maturation of the CPK in early cytoplasmic SSU precursors is not specific to yeast, but rather more generally conserved in eukaryotes. It still remains unclear how exactly the conversion of CPK-im to CPK is triggered. Direct interactions of rpS2/uS5 and helix h36 with helix h2 might be involved (see interactions of h2 in S7 Appendix, A), but also other, more indirect mechanisms which stimulate progressive maturation of the neck helix h28 seem plausible.

## Materials and Methods

### Yeast strains and microbiological procedures

Genome editing of yeast strains for expression of chromosomally encoded TAP-tagged versions of biogenesis factors was done using PCR amplified DNA fragments containing a TAP tag encoding element and a marker gene as described in [56]. For strains expressing the Enp1-TAP fusion protein, the PCR fragment was produced using the two oligonucleotides O2318 (5’-TTTGTT GATCCACAGGAAGCTAATGATGATTTAATGATTGATGTCAATTCCATGGAAAAGAGAAG-3’) and O2319 (5’-GGGAAAGACCGAGCGATATAAAATTGATGAAAAATTGATATTACAGCATACGA CTCACTATAGGG-3’) together with the template plasmid pBS1539 [56]. Yeast strains BY4741 (Euroscarf), Y801, and Y1236 (both [33]) were transformed with the resulting fragment. For the strain expressing Slx9-TAP, a PCR fragment was produced using the two oligonucleotides O3877 (5’-AATCCATTTGGCGCCTTAAGAGAGGTTATCAAGCTGCAAAAACAATCCATGGAAAAG AGAAG-3’) and O3878 (5’-TATATATTACACTGGCAAAAATTGTTATGCTATGCTATTTAATGTTACACTCACTATAGGG-3’) together with plasmid pBS1539 as a template. Here, yeast strain Y801 [33] was transformed with the resulting fragment. After transformation [57] and selection on plates lacking uracil, clones were further analysed for expression of the fusion protein by western blotting to obtain yeast strains Y2109 (his3-1, leu2-0, ura3-0, RPS21A::kanMX4, RPS21B::kanMX4, ENP1::TAP-klURA3), Y2110 (his3-1, leu2-0, ura3-0, RPS29B::kanMX4, RPS29A::HIS3MX6, ENP1::TAP-klURA3), Y2226 (his3-1, leu2-0, met15-0, ura3-0, ENP1::TAP-klURA3) and Y3153 (his3-1, leu2-0, ura3-0, RPS21A::kanMX4, RPS21B::kanMX4, SLX9::TAP-klURA3).

### Affinity purification of ribosomal particles and cryo-EM analyses

Cells of 8 liters of a culture of strains Y2226 and Y3153, and of 2l of a culture of strain Y2109 were used for extract preparation and affinity purification. After logarithmic growth in YPG medium (1% yeast extract, 2% bacto peptone, 2% galactose) all cultures were shifted for four hours at 30° C to YPD medium (1% yeast extract, 2% bacto peptone, 2% glucose) before harvesting the cells.Preparation of cellular extracts, one-step affinity-purification of Enp1-TAP or Slx9-TAP associated pre-ribosomal particles, and vitrification of samples on holey carbon R 1.2/1.3 copper 300 mesh grids (Quantifoil) using a Vitrobot MarkIV device (Thermofisher) were performed as described in [58]. Final data acquisition was done at 200 kV and at a magnification of 50k on a JEM-Z200FSC (JEOL) electron microscope equipped with a K2 Summit direct electron detector camera (Gatan). Images were acquired using the SerialEM software package [59,60]. The camera was used in counting mode, 40 frames were acquired per image movie, the pixel size was 0.968 Å/pixel, the total dose per movie 40 electrons/Å, and the total exposure time was 4.4 seconds. The target defocus range was set between −0.8 and −2.2 micrometers.

### RNA extraction and northern blotting

RNA extraction and northern blotting were performed as described in [58], using probes hybridizing in the ITS1 (O1819, 5’-GTAAAAGCTCTCATGCTCTTGCC-3’), in the 25S rRNA (O212, 5’-CTCCGCTTATTGATATGC-3’), or in the 18S rRNA (O205, 5’-CATGGCTTAATCTTTGAGAC-3’).Signals on image plates were quantified using the ImageJ software package.

### Cryo-EM data processing and structure model fitting, interpretation, and visualization

Cryo-EM data processing was performed in Relion 4.0 [61]. Beam-induced motion correction was done using Relions own implementation of the MotionCor2 algorithm [62]. Contrast transfer functions of images were estimated using CTFFIND4.1 [63]. Candidate particles were picked using the Topaz particle-picking pipeline with the provided pre-trained model [64]. More details on single particle processing and classification strategies for the different experimental datasets are shown in S2, S4, and S6 Appendices. Average map resolutions were estimated as implemented in Relion according to the Fourier shell correlation at 0.143 between two independently refined half-maps [65,66].

The Protein Data Bank entry 6EML was used as starting model [26] to build the Enp1TAP_A and Enp1TAP-S21_A models. For yeast Nob1 and parts of yeast Tsr1, their respective structure predictions by the AlphaFold algorithm were used as starting points for their model building [53,54]. Initial rigid body fitting and model editing were done in UCSF ChimeraX [67] and in Coot [68]. The fit of the models to the respective experimental maps was further improved by molecular dynamics flexible fitting using the Isolde plugin in UCSF ChimeraX [69]. Fitting was first done for the rRNA and its surrounding with distance restraints applied between nucleobases (typically distance cutoff 5 and kappa 200) which were then gradually relaxed. Subsequently, individual protein chains were flexibly fitted. The fit and geometry of each RNA and protein chain were then inspected and if necessary manually corrected for individual residues, or in part newly built (Nob1 C-terminus), using the Isolde plugin in ChimeraX. Final model refinement was done using the phenix.real_space refinement tool [70] with starting parameters created from the Isolde plugin by the command ‘isolde write phenixRsrInput’ after a global molecular dynamics simulation. Model statistics shown in S5 and S7 Appendixes were obtained in UCSF ChimeraX using a python script in which possible direct residue interactions were predicted by the ChimeraX command ‘contacts’ with default settings for ‘overlapCutoff’ and ‘hbondAllowance’ parameters. Average map values of proteins were determined by applying the ChimeraX ‘fitmap’ command for individual chains in respective maps. Yeast LSU rRNA domain and helix definitions were taken from [3]. The model of the mature small ribosomal subunit represented in Fig 2 is protein database entry 4v88 [2]. All representations of structure models or maps were created in ChimeraX.

### Scoring approach to identify candidate 2’-O-methyl adenosine residues

A three-step approach was used to rank adenosines in the given models and maps for possible methyl modification at the 2’-O atom. Firstly, all adenosine residues in the examined models were converted into 2’-O-methyl adenosine residues by a python script in the ChimeraX (1.4) environment using the ‘build modify’ command. The torsion of the bond between the 2’O and the newly created methyl group was adapted for each residue to avoid clashes in the model. In a second step, a molecular dynamics simulation was performed in ChimeraX using the Isolde plugin with the modified model and the respective map. The simulation was set to ‘High Fidelity’ and the map weight to 0.19. The forcefield used for the 2’-O-methyl adenosine residue was adapted from the RNA-OL3 nucleic acid force field definitions [71]. In the third step, scoring was performed using another python script in the ChimeraX environment. Here, a first map value was determined at each adenosine for the 2’-O-methyl atom using the ‘fit_atoms_in_map” function. Subsequently, a second map value was determined at each adenosine for the 2’-O-methyl atom after prolonging the bond between the 2’O atom and the 2’-O-methyl atom to 2.4 Å. The first map value was used to determine the score ‘Sc_MapVal’ for each adenosine, and the score ‘SC_FallOff’ was calculated for each adenosine by dividing the first by the second map value.

## Supporting information

S1 Raw Images

## Acknowledgments

We are grateful to Dr. Michael Pilsl for his support during cryo-EM data acquisition and Dr. Christoph Engel for the critical inspection of de novo modelled parts of the structure models (both University of Regensburg, Regensburg Center of Biochemistry, Structural Biochemistry). We thank Dr Sébastien Ferreira-Cerca and Michel Jüttner for sharing their expertise on SSU maturation in prokaryotic organisms (both University of Regensburg, Regensburg Center of Biochemistry, Cellular Biochemistry of Microorganisms). We thank Drs. Steffen Jakob and Thomas Hierlmeier for the construction of yeast strains used in this work, and former and present members of the chairs for biochemistry I and III for their constant support and for helpful discussions.

## Supporting Information captions

**S1 Appendix. Analysis of the (pre-)rRNA composition of Enp1-TAP or Slx9-TAP associated particles purified from yeast r-protein expression mutants or a control strain**.

Cells of yeast strains Y2226, Y2109, Y2110, and Y3153 which express the indicated r-protein (‘depleted’) under control of the GAL1/10 promoter, and the indicated biogenesis factor (‘Bait’) in fusion with the TAP-tag were incubated for four hours in a glucose-containing medium (see Materials and Methods). Corresponding cellular extracts were used for affinity purification of the bait proteins and (pre-)rRNA composition of the extracts (‘Input’) and final eluates (‘Eluate’) were analyzed by RNA extraction and northern blotting with probes O1819 (A) and a mixture of probes O205 and O212 (B). On the left of (A) and (B), the detected (pre-)rRNA species are designated. The ratios indicated in (C) of the 20S pre-rRNA to 18S rRNA and of the 25S rRNA to 18S rRNA ratio in the input fractions were determined with ImageJ. The conditional expression mutant of the S3-cluster r-protein rpS29/uS14 (lanes 3 and 7) was included for comparison.

**S2 Appendix. Single particle sorting and processing scheme for the cryo-EM dataset of particles purified via Enp1-TAP from control cells (strain Y2226)**.

**S3 Appendix. Summary of 2’-O-methyl scoring for Enp1TAP_A (A) and Enp1TAP-S21_A (B) maps and models**.

**S4 Appendix. Single particle sorting and processing scheme for the cryo-EM dataset of particles purified via Enp1-TAP from the rpS21/eS21 expression mutant strain Y2109**.

**S5 Appendix. Interactions of proteins modeled in Enp1TAP_A and Enp1TAP-S21_A**.

**S6 Appendix. Single particle sorting and processing scheme for the cryo-EM dataset of particles purified via Slx9-TAP from the rpS21/eS21 expression mutant strain Y3153**.

**S7 Appendix. Interactions of RNA helices modeled in Enp1TAP_A and Enp1TAP-S21_A**.

**S8 Appendix. Raw images**.

## Notes

### Competing Interest Statement

The authors have declared no competing interest.

