## Supplementary figures and images for "Impact of the yeast S0/uS2-cluster ribosomal protein rpS21/eS21 on rRNA folding and the architecture of small ribosomal subunit precursors"

### S1 Appendix

A B

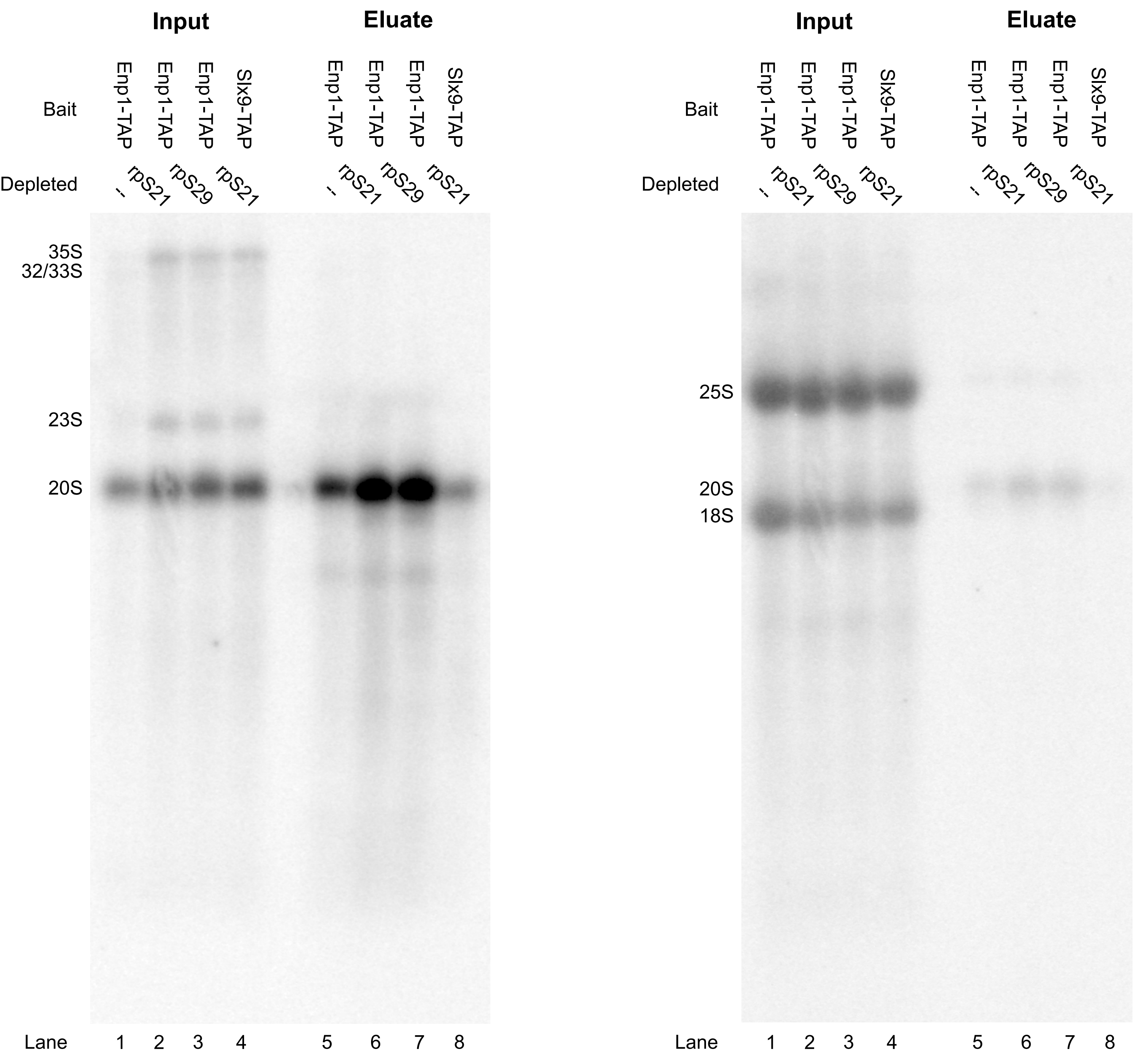


C

| rRNA in | Normalized ratio 20S:18S | Normalized ratio 25S:18S |
|---|---|---|
| **Lane 1** | 1,00 | 1,00 |
| **Lane 2** | 1,62 | 1,31 |
| **Lane 3** | 2,20 | 1,44 |
| **Lane 4** | 2,09 | 1,24 |

### S2 Appendix

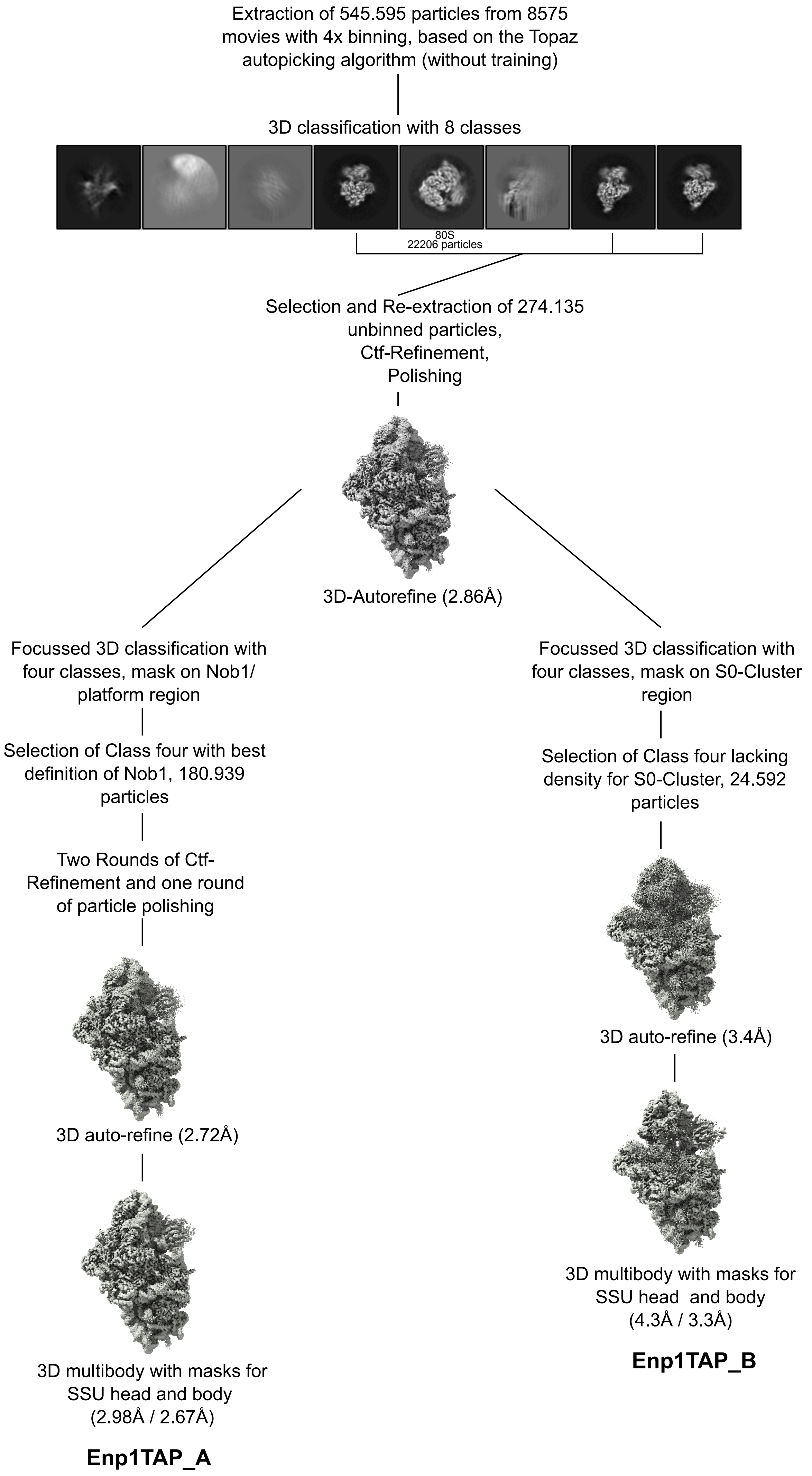

### S4 Appendix

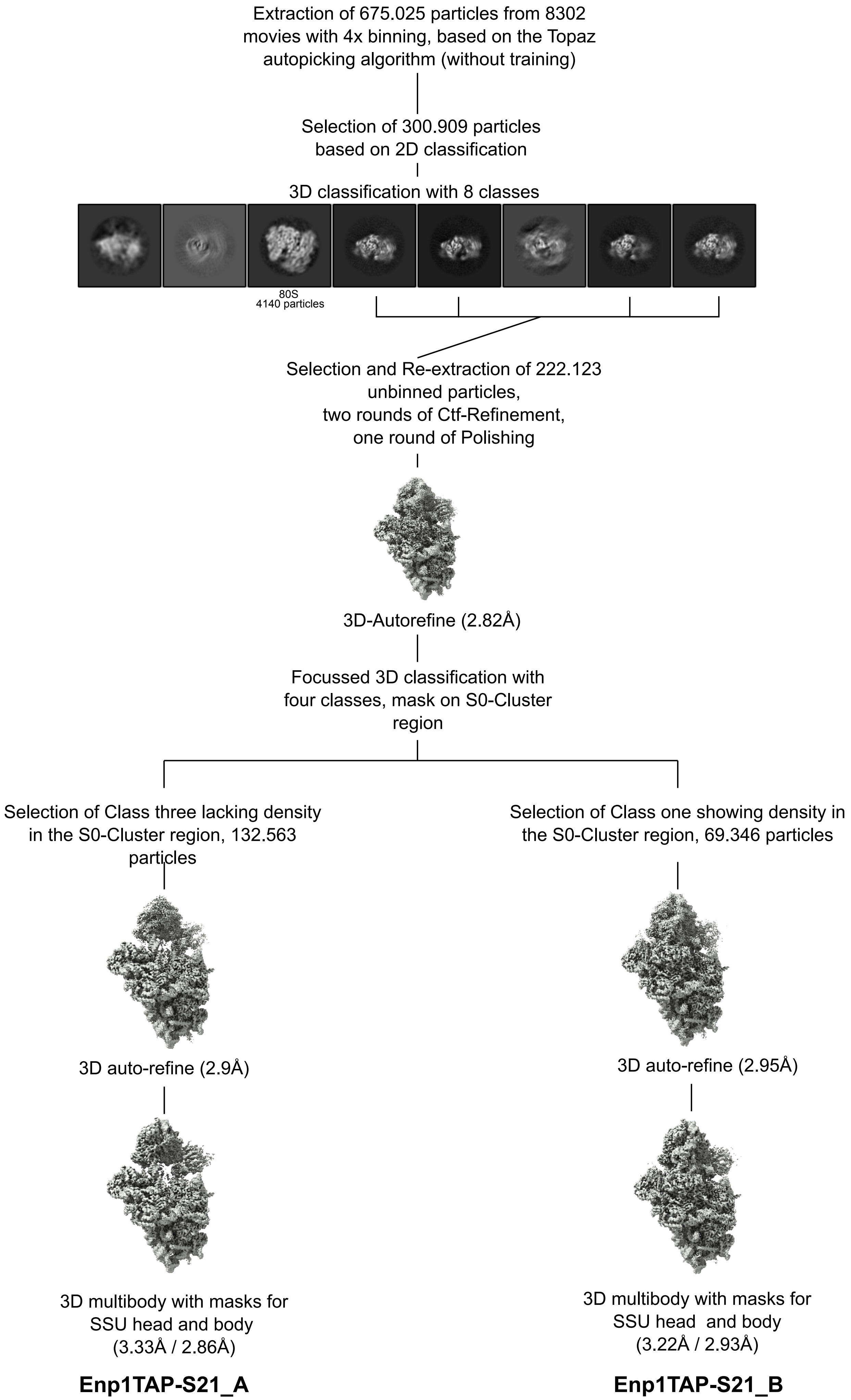
