## Supplementary material for "Impact of the yeast S0/uS2-cluster ribosomal protein rpS21/eS21 on rRNA folding and the architecture of small ribosomal subunit precursors": S3 Appendix

A)

| Position | Known AM modification | Sc_MapVal | Sc_FallOff |
|---|---|---|---|
| 1300 | no | 0,0158 | 1,82 |
| **796** | **yes** | **0,0147** | **2,10** |
| **100** | **yes** | **0,0146** | **2,02** |
| **541** | **yes** | **0,0135** | **1,71** |
| **28** | **yes** | **0,0134** | **1,88** |
| **974** | **yes** | **0,0131** | **1,67** |
| **619** | **yes** | **0,0125** | **2,30** |
| 752 | no | 0,0124 | 1,05 |
| 385 | no | 0,0112 | 0,83 |
| 407 | no | 0,0109 | 1,50 |
| 1754 | no | 0,0108 | 0,94 |
| 1391 | no | 0,0107 | 0,99 |
| **420** | **yes** | **0,0105** | **2,10** |
| 86 | no | 0,0104 | 1,13 |
| **436** | **yes** | **0,0102** | **3,56** |
| 1337 | no | 0,0102 | 1,31 |
| 579 | no | 0,0097 | 1,04 |
| 22 | no | 0,0097 | 1,21 |
| 599 | no | 0,0095 | 1,15 |
| 1183 | no | 0,0095 | 0,80 |
| 1559 | no | 0,0092 | 1,20 |
| 1384 | no | 0,0091 | 1,16 |
| 219 | no | 0,0091 | 1,01 |
| 1326 | no | 0,0091 | 1,18 |
| 1093 | no | 0,0090 | 1,01 |
| 1344 | no | 0,0089 | 1,21 |
| 844 | no | 0,0088 | 1,19 |
| 630 | no | 0,0088 | 1,59 |
| 46 | no | 0,0088 | 1,34 |
| 855 | no | 0,0087 | 1,32 |
| 399 | no | 0,0086 | 1,04 |
| 1689 | no | 0,0086 | 1,07 |
| 570 | no | 0,0085 | 1,29 |
| 606 | no | 0,0085 | 0,70 |
| 1444 | no | 0,0084 | 0,94 |
| 1411 | no | 0,0084 | 1,17 |
| 1025 | no | 0,0083 | 1,19 |
| 85 | no | 0,0083 | 0,79 |
| 585 | no | 0,0083 | 1,24 |
| 333 | no | 0,0082 | 1,30 |
| 1023 | no | 0,0082 | 1,34 |
| 1781 | no | 0,0082 | 1,21 |
| 505 | no | 0,0082 | 1,23 |
| 474 | no | 0,0082 | 1,10 |
| 1084 | no | 0,0082 | 1,10 |
| 370 | no | 0,0081 | 1,76 |
| 829 | no | 0,0081 | 1,20 |
| 416 | no | 0,0080 | 0,99 |
| 93 | no | 0,0080 | 1,27 |
| 515 | no | 0,0080 | 1,04 |
| 1313 | no | 0,0079 | 1,13 |
| 850 | no | 0,0079 | 1,35 |
| 72 | no | 0,0078 | 1,58 |
| 1139 | no | 0,0077 | 1,04 |
| 353 | no | 0,0077 | 1,24 |
| 43 | no | 0,0077 | 1,91 |
| 940 | no | 0,0077 | 1,01 |
| 62 | no | 0,0076 | 1,33 |
| 391 | no | 0,0076 | 3,56 |
| 1171 | no | 0,0076 | 1,08 |
| 331 | no | 0,0076 | 1,64 |

B)

| Position | Known AM modification | Sc_MapVal | Sc_FallOff |
|---|---|---|---|
| **100** | **yes** | **0,0147** | **1,70** |
| **796** | **yes** | **0,0140** | **2,39** |
| **28** | **yes** | **0,0139** | **1,84** |
| 752 | no | 0,0136 | 0,99 |
| 538 | no | 0,0134 | 2,05 |
| **974** | **yes** | **0,0130** | **1,68** |
| 1183 | no | 0,0129 | 1,06 |
| 86 | no | 0,0123 | 1,67 |
| 385 | no | 0,0121 | 0,83 |
| 407 | no | 0,0120 | 1,99 |
| **420** | **yes** | **0,0114** | **1,78** |
| 1142 | no | 0,0112 | 1,04 |
| 1753 | no | 0,0108 | 1,16 |
| **436** | **yes** | **0,0106** | **1,97** |
| 399 | no | 0,0106 | 1,61 |
| 456 | no | 0,0104 | 1,45 |
| 1714 | no | 0,0102 | 1,18 |
| **541** | **yes** | **0,0102** | **1,71** |
| 1093 | no | 0,0101 | 1,18 |
| 416 | no | 0,0101 | 0,96 |
| 1147 | no | 0,0099 | 1,09 |
| 360 | no | 0,0097 | 1,05 |
| 599 | no | 0,0097 | 1,15 |
| 967 | no | 0,0096 | 1,46 |
| 219 | no | 0,0096 | 0,86 |
| 1555 | no | 0,0095 | 1,39 |
| 1592 | no | 0,0095 | 1,00 |
| 46 | no | 0,0094 | 2,26 |
| 1689 | no | 0,0094 | 1,03 |
| 369 | no | 0,0094 | 0,96 |
| 1559 | no | 0,0092 | 1,12 |
| 770 | no | 0,0091 | 1,36 |
| 1163 | no | 0,0091 | 0,83 |
| 1746 | no | 0,0091 | 1,31 |
| 251 | no | 0,0091 | 1,79 |
| 1750 | no | 0,0091 | 1,22 |
| 585 | no | 0,0090 | 1,29 |
| 859 | no | 0,0090 | 1,25 |
| 847 | no | 0,0090 | 1,05 |
| 162 | no | 0,0089 | 1,13 |
| 323 | no | 0,0089 | 1,32 |
| 923 | no | 0,0088 | 1,24 |
| 518 | no | 0,0088 | 1,65 |
| **619** | **yes** | **0,0087** | **2,07** |
| 1492 | no | 0,0087 | 1,12 |
| 505 | no | 0,0086 | 1,17 |
| 391 | no | 0,0086 | 0,98 |
| 1730 | no | 0,0086 | 1,47 |
| 924 | no | 0,0086 | 1,54 |
| 353 | no | 0,0085 | 1,42 |
| 76 | no | 0,0085 | 1,12 |
| 441 | no | 0,0085 | 1,38 |
| 963 | no | 0,0085 | 0,82 |
| 485 | no | 0,0084 | 1,19 |
| 591 | no | 0,0084 | 2,02 |
| 301 | no | 0,0084 | 1,46 |
| 580 | no | 0,0084 | 0,98 |
| 217 | no | 0,0084 | 1,67 |
| 55 | no | 0,0083 | 1,76 |
| 829 | no | 0,0083 | 1,12 |
| 425 | no | 0,0083 | 0,97 |

A)

| Position | Known AM modification | Sc_MapVal | Sc_FallOff |
|---|---|---|---|
| 253 | no | 0,0076 | 1,60 |
| 1322 | no | 0,0076 | 1,28 |
| 76 | no | 0,0075 | 1,42 |
| 550 | no | 0,0075 | 1,71 |
| 814 | no | 0,0075 | 1,39 |
| 635 | no | 0,0075 | 1,36 |
| 1142 | no | 0,0075 | 3,96 |
| 438 | no | 0,0074 | 1,91 |
| 168 | no | 0,0074 | 1,09 |
| 1036 | no | 0,0074 | 1,14 |
| 1230 | no | 0,0074 | 1,08 |
| 256 | no | 0,0073 | 1,03 |
| 556 | no | 0,0073 | 0,98 |
| 456 | no | 0,0073 | 1,29 |
| 1750 | no | 0,0073 | 1,27 |
| 770 | no | 0,0073 | 1,00 |
| 998 | no | 0,0073 | 1,20 |
| 527 | no | 0,0072 | 1,36 |
| 1224 | no | 0,0072 | 1,33 |
| 1027 | no | 0,0072 | 1,54 |
| 1030 | no | 0,0072 | 1,54 |
| 545 | no | 0,0072 | 1,16 |
| 1492 | no | 0,0072 | 1,05 |
| 428 | no | 0,0072 | 2,09 |
| 971 | no | 0,0072 | 1,69 |
| 1147 | no | 0,0072 | 1,35 |
| 580 | no | 0,0071 | 1,16 |
| 1329 | no | 0,0071 | 0,93 |
| 251 | no | 0,0071 | 2,13 |
| 464 | no | 0,0071 | 1,34 |
| 47 | no | 0,0071 | 1,06 |
| 906 | no | 0,0071 | 1,34 |
| 1043 | no | 0,0071 | 1,49 |
| 1081 | no | 0,0071 | 1,07 |
| 1479 | no | 0,0071 | 1,40 |
| 1791 | no | 0,0071 | 1,24 |
| 1417 | no | 0,0071 | 1,12 |
| 1555 | no | 0,0071 | 1,61 |
| 1088 | no | 0,0071 | 1,71 |
| 266 | no | 0,0070 | 1,52 |
| 907 | no | 0,0070 | 1,12 |
| 905 | no | 0,0070 | 1,00 |
| 1600 | no | 0,0070 | 1,24 |
| 41 | no | 0,0070 | 0,96 |
| 1714 | no | 0,0070 | 1,46 |
| 1728 | no | 0,0070 | 1,78 |
| 425 | no | 0,0070 | 1,22 |
| 520 | no | 0,0070 | 1,18 |
| 1076 | no | 0,0069 | 1,51 |
| 1746 | no | 0,0069 | 1,66 |
| 11 | no | 0,0069 | 1,44 |
| 19 | no | 0,0069 | 1,46 |
| 526 | no | 0,0069 | 1,44 |
| 1446 | no | 0,0069 | 1,24 |
| 80 | no | 0,0069 | 1,23 |
| 601 | no | 0,0069 | 1,19 |
| 1202 | no | 0,0069 | 1,03 |
| 525 | no | 0,0069 | 0,97 |
| 847 | no | 0,0069 | 1,50 |
| 295 | no | 0,0068 | 1,33 |
| 301 | no | 0,0068 | 1,15 |

B)

| Position | Known AM modification | Sc_MapVal | Sc_FallOff |
|---|---|---|---|
| 844 | no | 0,0082 | 1,20 |
| 1671 | no | 0,0082 | 1,50 |
| 93 | no | 0,0082 | 1,34 |
| 606 | no | 0,0082 | 0,94 |
| 474 | no | 0,0081 | 1,17 |
| 119 | no | 0,0081 | 1,12 |
| 771 | no | 0,0081 | 1,99 |
| 1587 | no | 0,0080 | 1,32 |
| 473 | no | 0,0080 | 1,31 |
| 1781 | no | 0,0080 | 1,33 |
| 1043 | no | 0,0080 | 1,10 |
| 535 | no | 0,0079 | 1,38 |
| 72 | no | 0,0079 | 1,31 |
| 906 | no | 0,0079 | 1,05 |
| 1691 | no | 0,0079 | 1,09 |
| 1160 | no | 0,0079 | 1,24 |
| 525 | no | 0,0078 | 1,09 |
| 1171 | no | 0,0078 | 0,95 |
| 570 | no | 0,0078 | 1,27 |
| 200 | no | 0,0078 | 1,55 |
| 1069 | no | 0,0078 | 1,61 |
| 1791 | no | 0,0077 | 1,05 |
| 978 | no | 0,0077 | 0,95 |
| 271 | no | 0,0077 | 1,19 |
| 218 | no | 0,0076 | 1,34 |
| 515 | no | 0,0076 | 1,37 |
| 1586 | no | 0,0076 | 1,10 |
| 80 | no | 0,0076 | 1,26 |
| 19 | no | 0,0076 | 1,36 |
| 1648 | no | 0,0076 | 1,30 |
| 1184 | no | 0,0076 | 1,44 |
| 1357 | no | 0,0076 | 1,00 |
| 1776 | no | 0,0076 | 1,86 |
| 254 | no | 0,0076 | 1,40 |
| 438 | no | 0,0075 | 2,36 |
| 428 | no | 0,0075 | 1,39 |
| 621 | no | 0,0075 | 0,99 |
| 1782 | no | 0,0075 | 1,13 |
| 483 | no | 0,0075 | 1,22 |
| 173 | no | 0,0075 | 1,14 |
| 256 | no | 0,0075 | 1,23 |
| 446 | no | 0,0075 | 1,16 |
| 112 | no | 0,0074 | 1,43 |
| 793 | no | 0,0074 | 1,23 |
| 1025 | no | 0,0074 | 0,95 |
| 1471 | no | 0,0074 | 1,48 |
| 481 | no | 0,0074 | 1,42 |
| 550 | no | 0,0073 | 1,79 |
| 1651 | no | 0,0073 | 1,59 |
| 470 | no | 0,0073 | 1,46 |
| 1143 | no | 0,0073 | 1,43 |
| 85 | no | 0,0073 | 5,21 |
| 545 | no | 0,0073 | 1,49 |
| 148 | no | 0,0073 | 1,98 |
| 1234 | no | 0,0073 | 1,01 |
| 1113 | no | 0,0073 | 1,04 |
| 1152 | no | 0,0073 | 1,24 |
| 344 | no | 0,0072 | 1,35 |
| 630 | no | 0,0072 | 1,37 |
| 979 | no | 0,0072 | 1,28 |
| 1139 | no | 0,0072 | 1,14 |

**A)**

| Position | Known AM modification | Sc_MapVal | Sc_FallOff |
|---|---|---|---|
| 1061 | no | 0,0068 | 1,64 |
| 156 | no | 0,0068 | 1,31 |
| 791 | no | 0,0068 | 1,18 |
| 1592 | no | 0,0068 | 1,21 |
| 892 | no | 0,0068 | 1,13 |
| 475 | no | 0,0067 | 1,17 |
| 1375 | no | 0,0067 | 1,30 |
| 481 | no | 0,0067 | 1,44 |
| 623 | no | 0,0067 | 1,98 |
| 247 | no | 0,0067 | 1,21 |
| 483 | no | 0,0067 | 1,18 |
| 220 | no | 0,0067 | 1,17 |
| 592 | no | 0,0066 | 2,06 |
| 315 | no | 0,0066 | 1,24 |
| 518 | no | 0,0066 | 1,47 |
| 898 | no | 0,0066 | 0,87 |
| 1223 | no | 0,0066 | 1,19 |
| 930 | no | 0,0066 | 1,43 |
| 55 | no | 0,0066 | 0,83 |
| 1086 | no | 0,0066 | 1,16 |
| 1631 | no | 0,0066 | 0,95 |
| 210 | no | 0,0066 | 1,48 |
| 182 | no | 0,0066 | 1,55 |
| 473 | no | 0,0065 | 1,07 |
| 1659 | no | 0,0065 | 1,18 |
| 756 | no | 0,0065 | 1,40 |
| 1556 | no | 0,0065 | 0,86 |
| 1691 | no | 0,0065 | 1,10 |
| 859 | no | 0,0065 | 1,16 |
| 359 | no | 0,0064 | 1,31 |
| 1794 | no | 0,0064 | 1,59 |
| 344 | no | 0,0064 | 1,20 |
| 762 | no | 0,0064 | 1,30 |
| 1776 | no | 0,0064 | 3,59 |
| 622 | no | 0,0064 | 1,29 |
| 400 | no | 0,0063 | 0,80 |
| 1116 | no | 0,0063 | 1,51 |
| 1587 | no | 0,0063 | 5,15 |
| 812 | no | 0,0063 | 1,31 |
| 1583 | no | 0,0063 | 0,99 |
| 1655 | no | 0,0063 | 1,16 |
| 594 | no | 0,0063 | 1,23 |
| 162 | no | 0,0063 | 1,58 |
| 323 | no | 0,0063 | 1,21 |
| 417 | no | 0,0063 | 1,89 |
| 534 | no | 0,0063 | 1,40 |
| 451 | no | 0,0063 | 2,90 |
| 891 | no | 0,0063 | 1,19 |
| 1651 | no | 0,0063 | 1,26 |
| 26 | no | 0,0063 | 1,29 |
| 1062 | no | 0,0063 | 1,68 |
| 774 | no | 0,0062 | 1,36 |
| 1143 | no | 0,0062 | 0,91 |
| 119 | no | 0,0062 | 1,09 |
| 1782 | no | 0,0062 | 1,17 |
| 254 | no | 0,0062 | 1,63 |
| 1092 | no | 0,0062 | 1,32 |
| 145 | no | 0,0062 | 1,21 |
| 299 | no | 0,0062 | 1,88 |
| 856 | no | 0,0062 | 1,38 |
| 1319 | no | 0,0062 | 1,31 |

**B)**

| Position | Known AM modification | Sc_MapVal | Sc_FallOff |
|---|---|---|---|
| 26 | no | 0,0072 | 1,46 |
| 295 | no | 0,0072 | 2,25 |
| 265 | no | 0,0072 | 1,07 |
| 1005 | no | 0,0072 | 1,02 |
| 71 | no | 0,0072 | 1,15 |
| 412 | no | 0,0072 | 1,14 |
| 966 | no | 0,0072 | 2,25 |
| 182 | no | 0,0071 | 1,33 |
| 1088 | no | 0,0071 | 2,33 |
| 1138 | no | 0,0071 | 1,81 |
| 65 | no | 0,0071 | 1,12 |
| 1081 | no | 0,0071 | 1,15 |
| 952 | no | 0,0071 | 1,45 |
| 1036 | no | 0,0071 | 1,43 |
| 520 | no | 0,0071 | 1,47 |
| 1667 | no | 0,0071 | 1,33 |
| 378 | no | 0,0071 | 2,21 |
| 512 | no | 0,0071 | 2,45 |
| 635 | no | 0,0070 | 1,16 |
| 1570 | no | 0,0070 | 1,37 |
| 812 | no | 0,0070 | 1,24 |
| 299 | no | 0,0070 | 2,30 |
| 437 | no | 0,0070 | 1,95 |
| 555 | no | 0,0070 | 1,55 |
| 817 | no | 0,0070 | 1,73 |
| 940 | no | 0,0070 | 1,08 |
| 534 | no | 0,0070 | 1,28 |
| 1211 | no | 0,0069 | 1,16 |
| 788 | no | 0,0069 | 1,08 |
| 1076 | no | 0,0069 | 1,20 |
| 1226 | no | 0,0069 | 1,15 |
| 755 | no | 0,0069 | 1,24 |
| 41 | no | 0,0069 | 1,02 |
| 47 | no | 0,0069 | 1,00 |
| 164 | no | 0,0069 | 1,70 |
| 1133 | no | 0,0069 | 1,14 |
| 198 | no | 0,0068 | 1,21 |
| 247 | no | 0,0068 | 4,28 |
| 1091 | no | 0,0068 | 1,32 |
| 1348 | no | 0,0068 | 1,22 |
| 22 | no | 0,0068 | 1,76 |
| 315 | no | 0,0068 | 0,97 |
| 971 | no | 0,0068 | 1,16 |
| 464 | no | 0,0068 | 1,06 |
| 61 | no | 0,0068 | 1,20 |
| 312 | no | 0,0068 | 1,10 |
| 1061 | no | 0,0068 | 1,82 |
| 1515 | no | 0,0068 | 0,85 |
| 1545 | no | 0,0068 | 1,61 |
| 147 | no | 0,0068 | 1,46 |
| 1556 | no | 0,0068 | 0,89 |
| 359 | no | 0,0068 | 1,17 |
| 526 | no | 0,0068 | 1,59 |
| 791 | no | 0,0068 | 1,38 |
| 811 | no | 0,0068 | 1,00 |
| 156 | no | 0,0067 | 1,22 |
| 884 | no | 0,0067 | 1,54 |
| 1593 | no | 0,0067 | 1,10 |
| 103 | no | 0,0067 | 0,91 |
| 370 | no | 0,0067 | 1,29 |
| 804 | no | 0,0067 | 1,61 |

A)

| Position | Known AM modification | Sc_MapVal | Sc_FallOff |
|---|---|---|---|
| 811 | no | 0,0062 | 1,10 |
| 615 | no | 0,0061 | 9,01 |
| 1570 | no | 0,0061 | 1,47 |
| 65 | no | 0,0061 | 1,16 |
| 112 | no | 0,0061 | 1,10 |
| 1388 | no | 0,0061 | 1,16 |
| 1019 | no | 0,0061 | 1,50 |
| 995 | no | 0,0061 | 1,26 |
| 1132 | no | 0,0061 | 1,44 |
| 1226 | no | 0,0061 | 0,96 |
| 1087 | no | 0,0061 | 2,08 |
| 1113 | no | 0,0061 | 1,97 |
| 1133 | no | 0,0061 | 1,33 |
| 1137 | no | 0,0061 | 1,35 |
| 221 | no | 0,0061 | 1,40 |
| 636 | no | 0,0061 | 1,05 |
| 471 | no | 0,0060 | -604,00 |
| 1157 | no | 0,0060 | 1,68 |
| 1204 | no | 0,0060 | 1,07 |
| 1020 | no | 0,0060 | 1,21 |
| 951 | no | 0,0060 | 1,31 |
| 955 | no | 0,0060 | 1,63 |
| 1160 | no | 0,0060 | 1,30 |
| 1345 | no | 0,0060 | 0,99 |
| 900 | no | 0,0060 | 1,01 |
| 1753 | no | 0,0060 | 1,17 |
| 1152 | no | 0,0060 | 1,21 |
| 244 | no | 0,0060 | 1,21 |
| 1515 | no | 0,0060 | 1,34 |
| 511 | no | 0,0059 | 2,18 |
| 591 | no | 0,0059 | 1,32 |
| 757 | no | 0,0059 | 2,84 |
| 485 | no | 0,0059 | 1,16 |
| 1712 | no | 0,0059 | 1,37 |
| 352 | no | 0,0059 | 0,98 |
| 988 | no | 0,0059 | 1,64 |
| 1184 | no | 0,0059 | 1,70 |
| 108 | no | 0,0059 | 1,39 |
| 441 | no | 0,0059 | 2,39 |
| 993 | no | 0,0059 | 1,19 |
| 1312 | no | 0,0059 | 1,35 |
| 206 | no | 0,0059 | 1,42 |
| 1005 | no | 0,0058 | 0,97 |
| 966 | no | 0,0058 | 1,90 |
| 1203 | no | 0,0058 | 1,28 |
| 535 | no | 0,0058 | 1,60 |
| 1221 | no | 0,0058 | 1,20 |
| 1211 | no | 0,0058 | 1,27 |
| 963 | no | 0,0058 | 0,74 |
| 126 | no | 0,0058 | 2,30 |
| 755 | no | 0,0058 | 1,16 |
| 983 | no | 0,0058 | 1,78 |
| 1296 | no | 0,0058 | 1,11 |
| 1721 | no | 0,0058 | 1,47 |
| 421 | no | 0,0057 | 1,45 |
| 923 | no | 0,0057 | 1,65 |
| 973 | no | 0,0057 | 2,06 |
| 1660 | no | 0,0057 | 1,30 |
| 788 | no | 0,0057 | 1,24 |
| 1238 | no | 0,0057 | 1,16 |
| 316 | no | 0,0057 | 1,35 |

B)

| Position | Known AM modification | Sc_MapVal | Sc_FallOff |
|---|---|---|---|
| 1487 | no | 0,0067 | 1,23 |
| 620 | no | 0,0067 | 5,05 |
| 998 | no | 0,0067 | 1,06 |
| 475 | no | 0,0067 | 1,37 |
| 1003 | no | 0,0067 | 1,09 |
| 1023 | no | 0,0067 | 1,34 |
| 257 | no | 0,0066 | 1,74 |
| 898 | no | 0,0066 | 0,97 |
| 907 | no | 0,0066 | 1,89 |
| 1483 | no | 0,0066 | 0,92 |
| 542 | no | 0,0066 | 1,21 |
| 221 | no | 0,0066 | 1,29 |
| 78 | no | 0,0066 | 1,47 |
| 105 | no | 0,0066 | 1,31 |
| 1202 | no | 0,0066 | 1,04 |
| 1020 | no | 0,0066 | 1,50 |
| 333 | no | 0,0065 | 0,92 |
| 789 | no | 0,0065 | 1,55 |
| 1728 | no | 0,0065 | 2,03 |
| 527 | no | 0,0065 | 1,72 |
| 623 | no | 0,0065 | 3,52 |
| 2 | no | 0,0065 | 0,87 |
| 809 | no | 0,0065 | 1,60 |
| 1125 | no | 0,0065 | 1,26 |
| 124 | no | 0,0065 | 1,28 |
| 926 | no | 0,0065 | 1,23 |
| 929 | no | 0,0065 | 1,20 |
| 753 | no | 0,0065 | 1,02 |
| 933 | no | 0,0065 | 1,95 |
| 1242 | no | 0,0065 | 1,12 |
| 1479 | no | 0,0064 | 1,17 |
| 1794 | no | 0,0064 | 1,29 |
| 1230 | no | 0,0064 | 1,07 |
| 1678 | no | 0,0064 | 1,43 |
| 352 | no | 0,0064 | 1,18 |
| 1655 | no | 0,0064 | 1,26 |
| 869 | no | 0,0064 | 1,32 |
| 1224 | no | 0,0064 | 1,16 |
| 1086 | no | 0,0064 | 1,34 |
| 145 | no | 0,0063 | 1,21 |
| 636 | no | 0,0063 | 0,99 |
| 855 | no | 0,0063 | 1,16 |
| 202 | no | 0,0063 | 1,20 |
| 684 | no | 0,0063 | 1,38 |
| 1027 | no | 0,0063 | 1,88 |
| 51 | no | 0,0063 | 1,29 |
| 410 | no | 0,0063 | 1,30 |
| 387 | no | 0,0063 | 1,60 |
| 760 | no | 0,0063 | 1,10 |
| 1087 | no | 0,0063 | 1,03 |
| 1600 | no | 0,0063 | 0,96 |
| 43 | no | 0,0063 | 1,09 |
| 1744 | no | 0,0063 | 1,34 |
| 244 | no | 0,0062 | 1,38 |
| 511 | no | 0,0062 | 1,98 |
| 1712 | no | 0,0062 | 1,26 |
| 11 | no | 0,0062 | 2,07 |
| 452 | no | 0,0062 | 2,86 |
| 601 | no | 0,0062 | 1,29 |
| 951 | no | 0,0062 | 1,50 |
| 171 | no | 0,0062 | 1,37 |

A)

| Position | Known AM modification | Sc_MapVal | Sc_FallOff |
|---|---|---|---|
| 1471 | no | 0,0057 | 1,23 |
| 67 | no | 0,0057 | 1,43 |
| 164 | no | 0,0057 | 1,40 |
| 200 | no | 0,0057 | 1,27 |
| 807 | no | 0,0057 | 1,29 |
| 809 | no | 0,0057 | 1,73 |
| 1039 | no | 0,0057 | 1,65 |
| 1800 | no | 0,0057 | 1,11 |
| 1586 | no | 0,0056 | 3,76 |
| 605 | no | 0,0056 | 0,83 |
| 1336 | no | 0,0056 | 1,15 |
| 1545 | no | 0,0056 | 1,59 |
| 387 | no | 0,0056 | -26,71 |
| 1648 | no | 0,0056 | 1,67 |
| 213 | no | 0,0056 | 1,08 |
| 468 | no | 0,0056 | 1,11 |
| 215 | no | 0,0056 | 1,08 |
| 265 | no | 0,0056 | 1,20 |
| 412 | no | 0,0056 | 1,05 |
| 1678 | no | 0,0056 | 1,19 |
| 157 | no | 0,0056 | 2,13 |
| 1577 | no | 0,0056 | 1,42 |
| 1801 | no | 0,0056 | 1,09 |
| 684 | no | 0,0056 | 1,43 |
| 1341 | no | 0,0056 | 1,41 |
| 1740 | no | 0,0056 | 1,20 |
| 378 | no | 0,0055 | -8,39 |
| 218 | no | 0,0055 | 1,28 |
| 1732 | no | 0,0055 | 2,01 |
| 197 | no | 0,0055 | 1,09 |
| 529 | no | 0,0055 | 0,94 |
| 1543 | no | 0,0055 | 1,33 |
| 754 | no | 0,0055 | 1,23 |
| 887 | no | 0,0055 | 1,56 |
| 1550 | no | 0,0055 | 1,77 |
| 933 | no | 0,0055 | 0,96 |
| 806 | no | 0,0055 | 1,31 |
| 924 | no | 0,0055 | 2,93 |
| 979 | no | 0,0054 | 2,03 |
| 1547 | no | 0,0054 | 1,63 |
| 217 | no | 0,0054 | 0,97 |
| 369 | no | 0,0054 | 1,11 |
| 926 | no | 0,0054 | 1,03 |
| 140 | no | 0,0054 | 0,89 |
| 1227 | no | 0,0054 | 1,02 |
| 105 | no | 0,0054 | 1,26 |
| 544 | no | 0,0054 | 1,13 |
| 61 | no | 0,0054 | 1,35 |
| 445 | no | 0,0054 | 1,74 |
| 799 | no | 0,0054 | 1,55 |
| 789 | no | 0,0054 | 1,35 |
| 869 | no | 0,0054 | 1,60 |
| 881 | no | 0,0054 | 1,68 |
| 753 | no | 0,0054 | 0,92 |
| 1667 | no | 0,0053 | 1,38 |
| 1671 | no | 0,0053 | 1,04 |
| 103 | no | 0,0053 | 0,99 |
| 271 | no | 0,0053 | 1,53 |
| 437 | no | 0,0053 | 2,21 |
| 804 | no | 0,0053 | 1,31 |
| 179 | no | 0,0053 | 1,76 |

B)

| Position | Known AM modification | Sc_MapVal | Sc_FallOff |
|---|---|---|---|
| 331 | no | 0,0062 | 1,56 |
| 451 | no | 0,0062 | 1,92 |
| 1092 | no | 0,0062 | 1,23 |
| 1719 | no | 0,0062 | 1,42 |
| 84 | no | 0,0061 | 1,40 |
| 1132 | no | 0,0061 | 1,57 |
| 988 | no | 0,0061 | 0,96 |
| 1475 | no | 0,0061 | 1,14 |
| 977 | no | 0,0061 | 1,29 |
| 213 | no | 0,0061 | 1,69 |
| 850 | no | 0,0061 | 1,30 |
| 400 | no | 0,0060 | 0,86 |
| 905 | no | 0,0060 | 1,23 |
| 1124 | no | 0,0060 | 1,35 |
| 529 | no | 0,0060 | 1,42 |
| 1740 | no | 0,0060 | 1,33 |
| 799 | no | 0,0060 | 2,13 |
| 1157 | no | 0,0060 | 1,16 |
| 754 | no | 0,0060 | 1,02 |
| 253 | no | 0,0060 | 1,24 |
| 1576 | no | 0,0059 | 1,24 |
| 992 | no | 0,0059 | 1,54 |
| 993 | no | 0,0059 | 1,07 |
| 222 | no | 0,0059 | 1,46 |
| 417 | no | 0,0059 | 1,25 |
| 1660 | no | 0,0059 | 1,83 |
| 973 | no | 0,0058 | 1,33 |
| 1244 | no | 0,0058 | 0,97 |
| 1493 | no | 0,0058 | 1,17 |
| 930 | no | 0,0058 | 1,15 |
| 685 | no | 0,0058 | 1,37 |
| 891 | no | 0,0058 | 1,37 |
| 104 | no | 0,0058 | 1,25 |
| 622 | no | 0,0058 | 1,15 |
| 92 | no | 0,0058 | 1,07 |
| 210 | no | 0,0058 | 10,87 |
| 605 | no | 0,0057 | 1,14 |
| 806 | no | 0,0057 | 1,51 |
| 915 | no | 0,0057 | 1,61 |
| 180 | no | 0,0057 | 1,57 |
| 1631 | no | 0,0057 | 1,13 |
| 62 | no | 0,0057 | 1,64 |
| 1208 | no | 0,0057 | 1,10 |
| 220 | no | 0,0056 | 1,24 |
| 126 | no | 0,0056 | 1,35 |
| 157 | no | 0,0056 | 1,14 |
| 604 | no | 0,0056 | 1,32 |
| 1797 | no | 0,0056 | 1,38 |
| 197 | no | 0,0056 | 1,02 |
| 881 | no | 0,0056 | 1,48 |
| 1203 | no | 0,0056 | 1,27 |
| 1227 | no | 0,0056 | 1,21 |
| 1611 | no | 0,0056 | 1,14 |
| 468 | no | 0,0056 | 0,98 |
| 1001 | no | 0,0056 | 0,96 |
| 892 | no | 0,0056 | 2,51 |
| 108 | no | 0,0055 | 1,12 |
| 544 | no | 0,0055 | 1,26 |
| 316 | no | 0,0055 | 1,26 |
| 1030 | no | 0,0055 | 1,64 |
| 1597 | no | 0,0055 | 1,31 |

A)

| Position | Known AM modification | Sc_MapVal | Sc_FallOff |
|---|---|---|---|
| 312 | no | 0,0053 | 1,31 |
| 1516 | no | 0,0053 | 1,05 |
| 1475 | no | 0,0053 | 0,98 |
| 567 | no | 0,0053 | 1,03 |
| 1124 | no | 0,0052 | 1,14 |
| 39 | no | 0,0052 | 0,89 |
| 181 | no | 0,0052 | 1,25 |
| 1573 | no | 0,0052 | 1,18 |
| 1208 | no | 0,0052 | 1,20 |
| 360 | no | 0,0052 | 0,70 |
| 771 | no | 0,0052 | 1,33 |
| 1348 | no | 0,0052 | 1,43 |
| 1242 | no | 0,0052 | 0,85 |
| 1013 | no | 0,0051 | 1,47 |
| 1360 | no | 0,0051 | 1,36 |
| 760 | no | 0,0051 | 1,48 |
| 929 | no | 0,0051 | 1,27 |
| 366 | no | 0,0051 | 1,12 |
| 51 | no | 0,0051 | 1,32 |
| 410 | no | 0,0051 | 1,54 |
| 939 | no | 0,0051 | 0,90 |
| 68 | no | 0,0051 | 1,27 |
| 1069 | no | 0,0051 | 2,65 |
| 1524 | no | 0,0051 | 7,57 |
| 171 | no | 0,0051 | 1,06 |
| 180 | no | 0,0051 | 1,35 |
| 685 | no | 0,0051 | 1,40 |
| 1526 | no | 0,0051 | 1,38 |
| 793 | no | 0,0050 | 0,94 |
| 173 | no | 0,0050 | 1,16 |
| 446 | no | 0,0050 | 1,76 |
| 1325 | no | 0,0050 | 1,28 |
| 542 | no | 0,0050 | 1,07 |
| 978 | no | 0,0050 | 0,97 |
| 555 | no | 0,0050 | 1,64 |
| 341 | no | 0,0050 | 0,86 |
| 1790 | no | 0,0050 | 3,76 |
| 769 | no | 0,0050 | 0,93 |
| 803 | no | 0,0050 | 1,09 |
| 1001 | no | 0,0049 | 1,40 |
| 538 | no | 0,0049 | 0,62 |
| 84 | no | 0,0049 | 1,29 |
| 40 | no | 0,0049 | 0,78 |
| 1460 | no | 0,0049 | 1,40 |
| 1576 | no | 0,0049 | 1,76 |
| 1166 | no | 0,0049 | 1,48 |
| 2 | no | 0,0048 | -2,63 |
| 148 | no | 0,0048 | 1,73 |
| 1505 | no | 0,0048 | 0,76 |
| 1125 | no | 0,0048 | 1,09 |
| 477 | no | 0,0048 | 2,14 |
| 884 | no | 0,0048 | 3,32 |
| 1219 | no | 0,0048 | 1,25 |
| 1483 | no | 0,0048 | 0,92 |
| 1091 | no | 0,0047 | 1,29 |
| 288 | no | 0,0047 | 0,98 |
| 1611 | no | 0,0047 | 1,67 |
| 1469 | no | 0,0047 | 1,52 |
| 862 | no | 0,0047 | 2,38 |
| 1321 | no | 0,0047 | 1,85 |
| 620 | no | 0,0047 | 1,00 |

B)

| Position | Known AM modification | Sc_MapVal | Sc_FallOff |
|---|---|---|---|
| 1732 | no | 0,0055 | 1,40 |
| 140 | no | 0,0055 | 1,02 |
| 1790 | no | 0,0055 | 11,64 |
| 1084 | no | 0,0055 | 1,06 |
| 1505 | no | 0,0055 | 0,91 |
| 1469 | no | 0,0055 | 1,36 |
| 1524 | no | 0,0055 | 1,91 |
| 181 | no | 0,0054 | 1,40 |
| 1062 | no | 0,0054 | 1,43 |
| 1681 | no | 0,0054 | 0,99 |
| 615 | no | 0,0053 | 1,19 |
| 1116 | no | 0,0053 | 1,22 |
| 215 | no | 0,0053 | 1,52 |
| 756 | no | 0,0053 | 1,49 |
| 1516 | no | 0,0053 | 0,95 |
| 807 | no | 0,0053 | 0,90 |
| 803 | no | 0,0053 | 0,98 |
| 814 | no | 0,0053 | 0,84 |
| 1221 | no | 0,0053 | 1,05 |
| 939 | no | 0,0052 | 1,39 |
| 1543 | no | 0,0052 | 1,09 |
| 1525 | no | 0,0052 | 1,28 |
| 288 | no | 0,0052 | 1,07 |
| 341 | no | 0,0052 | 1,08 |
| 421 | no | 0,0052 | 1,28 |
| 1569 | no | 0,0052 | 1,14 |
| 1526 | no | 0,0051 | 1,68 |
| 1375 | no | 0,0051 | 1,28 |
| 1659 | no | 0,0051 | 1,18 |
| 460 | no | 0,0050 | 10,44 |
| 521 | no | 0,0050 | -16,16 |
| 862 | no | 0,0050 | 1,36 |
| 68 | no | 0,0050 | 1,19 |
| 1137 | no | 0,0050 | 2,63 |
| 1223 | no | 0,0050 | 1,18 |
| 1550 | no | 0,0050 | 1,34 |
| 179 | no | 0,0049 | 1,27 |
| 900 | no | 0,0049 | 0,89 |
| 1360 | no | 0,0049 | 1,15 |
| 1749 | no | 0,0049 | 1,14 |
| 1039 | no | 0,0049 | 1,03 |
| 1721 | no | 0,0049 | 1,30 |
| 1238 | no | 0,0049 | 1,10 |
| 944 | no | 0,0048 | 0,99 |
| 1013 | no | 0,0048 | 1,01 |
| 367 | no | 0,0048 | 1,98 |
| 995 | no | 0,0048 | 1,54 |
| 266 | no | 0,0048 | 1,01 |
| 1547 | no | 0,0047 | 2,16 |
| 970 | no | 0,0047 | 0,99 |
| 1166 | no | 0,0047 | 1,06 |
| 1583 | no | 0,0047 | 1,00 |
| 366 | no | 0,0047 | 1,48 |
| 919 | no | 0,0046 | 2,62 |
| 1131 | no | 0,0046 | 0,96 |
| 567 | no | 0,0046 | 1,06 |
| 983 | no | 0,0046 | 1,14 |
| 1019 | no | 0,0045 | 1,26 |
| 67 | no | 0,0045 | 1,03 |
| 592 | no | 0,0045 | 1,21 |
| 556 | no | 0,0045 | 0,94 |

A)

| Position | Known AM modification | Sc_MapVal | Sc_FallOff |
|---|---|---|---|
| 71 | no | 0,0046 | 1,09 |
| 941 | no | 0,0046 | 2,76 |
| 521 | no | 0,0046 | 6,52 |
| 1487 | no | 0,0046 | 1,01 |
| 1797 | no | 0,0046 | 1,43 |
| 1410 | no | 0,0046 | 1,29 |
| 78 | no | 0,0046 | 2,95 |
| 621 | no | 0,0046 | 9,02 |
| 1236 | no | 0,0046 | 1,24 |
| 915 | no | 0,0046 | 1,46 |
| 1719 | no | 0,0046 | 1,03 |
| 1003 | no | 0,0045 | 1,12 |
| 257 | no | 0,0045 | 2,00 |
| 202 | no | 0,0045 | 1,05 |
| 124 | no | 0,0045 | 1,36 |
| 1722 | no | 0,0045 | 2,68 |
| 1593 | no | 0,0045 | 1,30 |
| 198 | no | 0,0045 | 1,21 |
| 1138 | no | 0,0044 | 1,77 |
| 977 | no | 0,0044 | 0,92 |
| 470 | no | 0,0044 | 1,07 |
| 944 | no | 0,0044 | 1,03 |
| 1597 | no | 0,0044 | 1,18 |
| 970 | no | 0,0044 | 1,00 |
| 1493 | no | 0,0044 | 1,25 |
| 147 | no | 0,0043 | 0,94 |
| 1382 | no | 0,0043 | 0,91 |
| 817 | no | 0,0043 | 1,59 |
| 1357 | no | 0,0043 | 2,13 |
| 92 | no | 0,0042 | 0,85 |
| 367 | no | 0,0041 | 3,76 |
| 1569 | no | 0,0041 | 1,18 |
| 746 | no | 0,0041 | 1,10 |
| 1749 | no | 0,0041 | 1,15 |
| 952 | no | 0,0041 | 0,88 |
| 919 | no | 0,0040 | 2,37 |
| 104 | no | 0,0040 | 1,14 |
| 1681 | no | 0,0040 | 0,90 |
| 1163 | no | 0,0040 | 1,67 |
| 452 | no | 0,0039 | 2,37 |
| 992 | no | 0,0039 | 1,12 |
| 1744 | no | 0,0039 | 1,35 |
| 1614 | no | 0,0037 | 0,83 |
| 1525 | no | 0,0037 | 0,86 |
| 1131 | no | 0,0035 | 1,35 |
| 328 | no | 0,0035 | -7,49 |
| 1503 | no | 0,0035 | 1,31 |
| 1802 | no | 0,0035 | 2,12 |
| 1065 | no | 0,0034 | 2,61 |
| 604 | no | 0,0033 | 0,71 |
| 865 | no | 0,0032 | 1,20 |
| 1256 | no | 0,0032 | 0,89 |
| 1234 | no | 0,0032 | 1,05 |
| 460 | no | 0,0032 | -3,42 |
| 967 | no | 0,0032 | -28,64 |
| 1731 | no | 0,0030 | -30,30 |
| 1026 | no | 0,0027 | -2,41 |
| 169 | no | 0,0026 | 0,49 |
| 512 | no | 0,0026 | 7,85 |
| 397 | no | 0,0026 | 1,06 |
| 300 | no | 0,0024 | 9,27 |

B)

| Position | Known AM modification | Sc_MapVal | Sc_FallOff |
|---|---|---|---|
| 887 | no | 0,0044 | 1,30 |
| 1204 | no | 0,0044 | 0,96 |
| 39 | no | 0,0044 | 1,13 |
| 168 | no | 0,0043 | 1,24 |
| 774 | no | 0,0043 | 1,32 |
| 1219 | no | 0,0043 | 2,00 |
| 1573 | no | 0,0043 | 1,10 |
| 477 | no | 0,0043 | 4,00 |
| 1026 | no | 0,0043 | -35,42 |
| 300 | no | 0,0042 | 9,77 |
| 445 | no | 0,0042 | 2,56 |
| 138 | no | 0,0041 | 0,87 |
| 1460 | no | 0,0041 | 1,41 |
| 1614 | no | 0,0040 | 1,28 |
| 865 | no | 0,0040 | 1,64 |
| 1256 | no | 0,0040 | 0,94 |
| 941 | no | 0,0040 | 1,44 |
| 40 | no | 0,0039 | 0,91 |
| 594 | no | 0,0039 | 0,96 |
| 206 | no | 0,0039 | 1,31 |
| 1236 | no | 0,0038 | 1,11 |
| 757 | no | 0,0038 | 1,54 |
| 955 | no | 0,0037 | 3,50 |
| 856 | no | 0,0036 | 1,82 |
| 1577 | no | 0,0036 | 1,15 |
| 169 | no | 0,0035 | 0,72 |
| 762 | no | 0,0035 | 1,12 |
| 746 | no | 0,0034 | 0,80 |
| 471 | no | 0,0034 | -7,28 |
| 769 | no | 0,0033 | 3,20 |
| 1722 | no | 0,0032 | 1,82 |
| 1503 | no | 0,0032 | 1,20 |
| 478 | no | 0,0032 | 5,89 |
| 1065 | no | 0,0031 | 2,15 |
| 401 | no | 0,0026 | 0,60 |
| 1731 | no | 0,0026 | -1,22 |
| 328 | no | 0,0025 | 25,10 |
| 863 | no | 0,0025 | 0,52 |
| 397 | no | 0,0017 | -5,03 |

A)

| Position | Known AM modification | Sc_MapVal | Sc_FallOff |
|---|---|---|---|
| 863 | no | 0,0020 | 0,71 |
| 1730 | no | 0,0019 | -2,77 |
| 401 | no | 0,0017 | 0,38 |
| 1244 | no | 0,0015 | 1,89 |
| 478 | no | 0,0015 | -10,36 |
| 138 | no | 0,0012 | 3,44 |

B)

| Position | Known AM modification | Sc_MapVal | Sc_FallOff |
|---|---|---|---|
