## Supplementary material for "Impact of the yeast S0/uS2-cluster ribosomal protein rpS21/eS21 on rRNA folding and the architecture of small ribosomal subunit precursors": S5 Appendix

| name | chain | modeled_residues Enp1TAP_A | mapped_contacts Enp1TAP_A | modeled_residues Enp1TAP-S21_A | mapped_contacts Enp1TAP-S21_A |
|---|---|---|---|---|---|
| rpS0 | P | 206 of 252 res.:<br>2-207; | rpS2(8);<br>rpS17(16);<br>RpS21(20);<br><br>Nob1(18);<br>H26(2);<br>H35(3);<br>H36(2);<br>H37(1); | - | - |
| rpS1 | Q | 214 of 255 res.:<br>20-233; | rpS14(5);<br><br>Pno1(2);<br>Nob1(2);<br>H22(5);<br>H23(5);<br>H23a(4);<br>H26(4);<br>H26_ES7(3); | 214 of 255 res.:<br>20-233; | rpS14(5);<br><br>H22(4);<br>H23(4);<br>H23a(4);<br>H26(3);<br>H26_ES7(3); |
| rpS2 | R | 207 of 254 res.:<br>35-87;<br>96-249; | rpS0(6);<br>rpS9(5);<br>rpS21(16);<br>rpS22(6);<br><br>H1(6);<br>H2(1);<br>H25(3);<br>H26a(2);<br>H35(1);<br>H36(2); | - | - |
| rpS4 | S | 260 of 261 res.:<br>2-261; | rpS6(6);<br>rpS9(4);<br>rpS24(8);<br><br>H6a(4);<br>H7(8);<br>H9(4);<br>H9_ES3b(1);<br>H12(3);<br>H13(4);<br>H15(5);<br>H21(8);<br>H21_unk(2);<br>H21_ES6a(2); | 260 of 261 res.:<br>2-261; | rpS6(5);<br>rpS9(5);<br>rpS24(7);<br><br>H6a(4);<br>H7(10);<br>H9(5);<br>H9_ES3b(1);<br>H12(2);<br>H13(3);<br>H15(5);<br>H21(9);<br>H21_unk(2);<br>H21_ES6a(2); |
| rpS5 | B | 201 of 225 res.:<br>20-149;<br>155-225; | rpS16(16);<br>rpS25(9);<br>rpS28(14);<br><br>H29(1);<br>H41(7);<br>H42(1);<br>H43(10); | 201 of 225 res.:<br>20-149;<br>155-225; | rpS16(15);<br>rpS25(9);<br>rpS28(13);<br><br>H29(2);<br>H41(7);<br>H42(2);<br>H43(10); |

| | | | | | |
|---|---|---|---|---|---|
| rpS6 | T | 226 of 236 res.:<br>1-226; | rpS4(4);<br>rpS24(1);<br><br>H6(9);<br>H7(2);<br>H8(22);<br>H9_ES3a(3);<br>H9(1);<br>H9_ES3b(2);<br>H10(6);<br>H13(5);<br>H14(3);<br>H44(4); | 226 of 236 res.:<br>1-226; | rpS4(4);<br><br>H6(10);<br>H7(2);<br>H8(22);<br>H9_ES3a(3);<br>H9(1);<br>H9_ES3b(2);<br>H10(6);<br>H13(5);<br>H14(3);<br>H44(3); |
| rpS7 | U | 184 of 190 res.:<br>4-187; | rpS13(2);<br>rpS22(8);<br><br>H20(2);<br>H21_ES6c(2);<br>H21_ES6b(2);<br>H21(4);<br>H21_ES6d(4);<br>H22(1); | 184 of 190 res.:<br>4-187; | rpS13(3);<br>rpS22(8);<br><br>H20(2);<br>H21_ES6c(3);<br>H21_ES6b(2);<br>H21(5);<br>H21_ES6d(5);<br>H22(1); |
| rpS8 | V | 185 of 200 res.:<br>2-122;<br>135-198; | rpS11(8);<br><br>H6a(2);<br>H7(4);<br>H9_ES3a(3);<br>H9(9);<br>H11(14);<br>H13(8);<br>H44(6); | 185 of 200 res.:<br>2-122;<br>135-198; | rpS11(10);<br><br>H6a(2);<br>H7(4);<br>H9_ES3a(2);<br>H9(9);<br>H11(15);<br>H13(8);<br>H44(6); |
| rpS9 | W | 185 of 197 res.:<br>2-186; | rpS2(5);<br>rpS4(4);<br>rpS24(2);<br>rpS30(5);<br><br>H1(2);<br>H3(1);<br>H4(8);<br>H12(2);<br>H16(2);<br>H17(8);<br>H18(4);<br>H21(4);<br>H21_unk(6); | 185 of 197 res.:<br>2-186; | rpS4(4);<br>rpS24(2);<br>rpS30(5);<br><br>H1(1);<br>H3(2);<br>H4(7);<br>H12(1);<br>H16(1);<br>H17(8);<br>H18(4);<br>H21(4);<br>H21_unk(6); |
| rpS11 | X | 141 of 156 res.:<br>5-145; | rpS8(9);<br>rpS22(1);<br>rpS23(8);<br><br>H7(6);<br>H9(6);<br>H11(14);<br>H12(1);<br>H19(2);<br>H20(1); | 141 of 156 res.:<br>5-145; | rpS8(15);<br>rpS22(2);<br>rpS23(8);<br><br>H7(5);<br>H9(5);<br>H11(14);<br>H12(1);<br>H19(2);<br>H20(1);<br>H21(1); |

| | | | | | |
|---|---|---|---|---|---|
| rpS13 | Y | 150 of 151 res.:<br>2-151; | rpS7(1);<br>rpS22(2);<br>rpS27(9);<br><br>H20(7);<br>H21_ES6d(2);<br>H22(21);<br>H23a(2);<br>H24(1);<br>H25(2);<br>H26(1); | 150 of 151 res.:<br>2-151; | rpS7(2);<br>rpS22(2);<br>rpS27(9);<br><br>H20(7);<br>H21_ES6d(2);<br>H22(21);<br>H23a(2);<br>H24(1);<br>H25(2); |
| rpS14 | Z | 127 of 137 res.:<br>11-137; | rpS1(8);<br>Pno1(16);<br><br>H23(17);<br>H23a(1);<br>H24(3);<br>H45(4); | 127 of 137 res.:<br>11-137; | rpS1(8);<br>Pno1(19);<br><br>H23(18);<br>H23a(1);<br>H24(3);<br>H45(4); |
| rpS15 | E | 114 of 142 res.:<br>14-69;<br>72-129; | rpS18(9);<br><br>H30(2);<br>H31_unk1(3);<br>H32(4);<br>H33(2);<br>H42(10); | 114 of 142 res.:<br>14-69;<br>72-129; | rpS18(11);<br><br>H30(2);<br>H31_unk1(4);<br>H32(5);<br>H33(1);<br>H42(10); |
| rpS16 | F | 125 of 143 res.:<br>3-127; | rpS5(19);<br>rpS19(1);<br><br>H38(2);<br>H39(2);<br>H39_ES9(2);<br>H41(5);<br>H43(6); | 125 of 143 res.:<br>3-127; | rpS5(20);<br><br>H39(1);<br>H39_ES9(2);<br>H41(5);<br>H43(4); |
| rpS17 | C | 61 of 136 res.:<br>65-125; | rpS0(18);<br>Nob1(1); | - | - |
| rpS18 | H | 122 of 146 res.:<br>10-131; | rpS15(11);<br>rpS19(5);<br>rpS25(2);<br><br>H41(3);<br>H42(10); | 122 of 146 res.:<br>10-131; | rpS15(12);<br>rpS19(4);<br>rpS25(1);<br><br>H41(3);<br>H42(10); |
| rpS19 | I | 141 of 144 res.:<br>3-143; | rpS16(1);<br>rpS18(6);<br><br>H30(3);<br>H39_ES9(7);<br>H41(18);<br>H42(3);<br>H43(2); | 141 of 144 res.:<br>3-143; | rpS18(5);<br><br>H30(3);<br>H39_ES9(7);<br>H41(20);<br>H42(2);<br>H43(2); |
| rpS21 | a | 86 of 87 res.:<br>1-86; | rpS0(16);<br>rpS2(17);<br>rpS22(2);<br>rpS27(5);<br><br>H26a(1); | - | - |

| | | | | | |
|---|---|---|---|---|---|
| rpS22 | b | 129 of 130 res.:<br>2-130; | rpS2(8);<br>rpS7(7);<br>rpS11(1);<br>rpS13(3);<br>rpS21(2);<br>rpS23(2);<br>rpS27(8);<br><br>H20(5);<br>H21_ES6c(2);<br>H21(8);<br>H22(3);<br>H25(11);<br>H26(1); | 129 of 130 res.:<br>2-130; | rpS7(6);<br>rpS11(2);<br>rpS13(3);<br>rpS23(1);<br>rpS27(6);<br><br>H20(5);<br>H21_ES6c(3);<br>H21(9);<br>H22(3);<br>H25(11);<br>H26(1); |
| rpS23 | c | 144 of 145 res.:<br>2-145; | rpS11(6);<br>rpS22(1);<br>Tsr1(11);<br><br>H3(6);<br>H4(1);<br>H5(1);<br>H11(4);<br>H12(1);<br>H15(1);<br>H18(9);<br>H19(5);<br>H20(1);<br>H25(6);<br>H27(2);<br>H44(1); | 144 of 145 res.:<br>2-145; | rpS11(6);<br>rpS22(1);<br>Tsr1(11);<br><br>H3(6);<br>H4(1);<br>H5(1);<br>H11(4);<br>H12(1);<br>H15(1);<br>H18(9);<br>H19(5);<br>H20(1);<br>H25(6);<br>H27(2);<br>H44(1); |
| rpS24 | d | 130 of 135 res.:<br>3-132; | rpS4(7);<br>rpS6(1);<br>rpS9(1);<br><br>H5(2);<br>H6(3);<br>H8(8);<br>H15(4);<br>H17(10);<br>H21_unk(1);<br>H21_ES6a(5); | 130 of 135 res.:<br>3-132; | rpS4(7);<br>rpS9(1);<br><br>H5(2);<br>H6(3);<br>H8(9);<br>H15(5);<br>H17(10);<br>H21_unk(1);<br>H21_ES6a(4); |
| rpS25 | K | 69 of 108 res.:<br>36-104; | rpS5(8);<br>rpS18(2);<br><br>H41(6); | 69 of 108 res.:<br>36-104; | rpS5(8);<br>rpS18(1);<br><br>H41(7); |
| rpS27 | f | 81 of 82 res.:<br>2-82; | rpS13(7);<br>rpS21(4);<br>rpS22(8);<br><br>H22(7);<br>H26(6); | 81 of 82 res.:<br>2-82; | rpS13(7);<br>rpS22(9);<br><br>H22(7);<br>H26(6); |
| rpS28 | L | 60 of 67 res.:<br>6-65; | rpS5(15);<br><br>H28(2);<br>H43(1); | 60 of 67 res.:<br>6-65; | rpS5(17);<br><br>H28(3);<br>H43(1); |

| | | | | | |
|---|---|---|---|---|---|
| rpS30 | g | 36 of 63 res.:<br>25-60; | rpS9(6);<br><br>H4(2);<br>H16(1);<br>H17(2);<br>H18(9); | 30 of 63 res.:<br>25-47;<br>54-60; | rpS9(6);<br><br>0(14);<br>H4(2);<br>H16(2);<br>H18(10); |
| Enp1 | e | 200 of 483 res.:<br>213-267;<br>273-326;<br>331-361;<br>367-379;<br>383-394;<br>414-427;<br>435-446;<br>454-462; | H32(2);<br>H34(1); | 179 of 483 res.:<br>213-267;<br>273-326;<br>331-361;<br>367-379;<br>383-394;<br>414-427; | H32(1);<br>H33(1); |
| Pno1 | p | 180 of 274 res.:<br>93-272; | rpS1(2);<br>rpS14(12);<br><br>Nob1(8);<br>H24(2);<br>H45(8); | 180 of 274 res.:<br>93-272; | rpS1(2);<br>rpS14(15);<br><br>H24(1);<br>H28(3);<br>H45(9); |
| Rio2 | r | 239 of 425 res.:<br>13-26;<br>31-42;<br>45-58;<br>77-92;<br>95-110;<br>120-126;<br>152-237;<br>247-287;<br>298-330; | Tsr1(1);<br><br>H29(3);<br>H30(3);<br>H42(2); | 190 of 425 res.:<br>78-92;<br>95-110;<br>120-126;<br>152-156;<br>158-237;<br>247-287;<br>298-313;<br>321-330; | H29(3);<br>H30(4);<br>H42(2); |
| Tsr1 | t | 598 of 788 res.:<br>44-114;<br>119-268;<br>273-309;<br>335-350;<br>463-786; | rpS23(7);<br>Rio2(1);<br><br>H5(5);<br>H7(1);<br>H11(3);<br>H15(1);<br>H17(2);<br>H27(2);<br>H31_unk1(3);<br>H31(2);<br>H32(2);<br>H34(1);<br>H44(6); | 619 of 788 res.:<br>12-32;<br>44-114;<br>119-268;<br>273-309;<br>335-350;<br>463-786; | rpS23(7);<br><br>H2(3);<br>H5(4);<br>H7(1);<br>H11(3);<br>H15(1);<br>H17(1);<br>H19(3);<br>H24(1);<br>H27(5);<br>H31_unk1(3);<br>H32(2);<br>H44(4);<br>H45(7); |
| Nob1 | o | 314 of 459 res.:<br>7-118;<br>221-249;<br>261-419;<br>444-457; | rpS0(18);<br>rpS1(3);<br>rpS17(1);<br>Pno1(11);<br><br>H23a(1);<br>H26(9);<br>H26a(2);<br>H35(1);<br>H36(4);<br>H37(4);<br>H45(1); | - | - |
