## Supplementary material for "Impact of the yeast S0/uS2-cluster ribosomal protein rpS21/eS21 on rRNA folding and the architecture of small ribosomal subunit precursors": S6 Appendix

Extraction of 318.619 particles from 6462 movies with 4x binning, based on the Topaz autopicking algorithm (without training)

3D classification with 8 classes

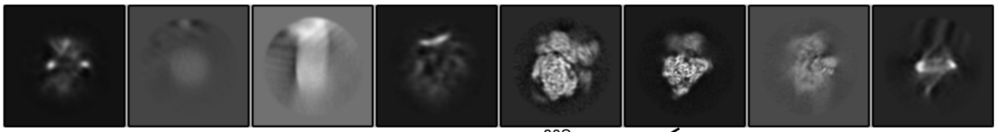

80S
10573 particles

Selection and Re-extraction of 53.304 unbinned particles,
one round of Ctf-Refinement,
one round of Polishing

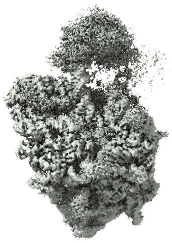

3D-Autorefine (3.3Å)

**Slx9TAP-S21_A**
