## Supplementary material for "Impact of the yeast S0/uS2-cluster ribosomal protein rpS21/eS21 on rRNA folding and the architecture of small ribosomal subunit precursors": S7 Appendix

| Helix | modeled_residues Enp1TAP_A | mapped_contacts Enp1TAP_A | modeled_residues Enp1TAP-S21_A | mapped_contacts Enp1TAP-S21_A |
|---|---|---|---|---|
| H1 | 13 of 14<br>2-8;<br>15-20; | rpS2(7);<br>rpS9(2);<br><br>H2(4);<br>H3(2);<br>H12(1);<br>H18(4);<br>H19(2);<br>H27(1); | 13 of 14<br>2-8;<br>15-20; | rpS9(1);<br><br>H2(4);<br>H3(2);<br>H12(1);<br>H18(3);<br>H19(2);<br>H27(1); |
| H2 | 14 of 14<br>9-14;<br>1137-1144; | rpS2(3);<br><br>H1(5);<br>H19(2);<br>H26a(3);<br>H27(1);<br>H28(1);<br>H36(4); | 14 of 14<br>9-14;<br>1137-1144; | Tsr1(2);<br><br>H1(5);<br>H19(2);<br>H26a(3);<br>H27(1);<br>H28(2); |
| H3 | 23 of 23<br>21-31;<br>595-606; | rpS9(1);<br>rpS23(8);<br><br>H1(1);<br>H4(2);<br>H12(4);<br>H15(1);<br>H18(3);<br>H19(1); | 23 of 23<br>21-31;<br>595-606; | rpS9(2);<br>rpS23(8);<br><br>H1(2);<br>H4(1);<br>H12(3);<br>H15(1);<br>H18(3); |
| H4 | 23 of 23<br>32-41;<br>466-478; | rpS9(14);<br>rpS23(1);<br>rpS30(6);<br><br>H3(1);<br>H15(3);<br>H17(6);<br>H18(2);<br>H21_unk(2); | 23 of 23<br>32-41;<br>466-478; | rpS9(10);<br>rpS23(1);<br>rpS30(6);<br><br>H3(1);<br>H15(3);<br>H17(5);<br>H18(3);<br>H21_unk(2); |
| H5 | 25 of 25<br>42-55;<br>424-434; | rpS23(1);<br>rpS24(3);<br>Tsr1(4);<br><br>H6a(5);<br>H7(1);<br>H12(1);<br>H13(2);<br>H15(6); | 25 of 25<br>42-55;<br>424-434; | rpS23(1);<br>rpS24(3);<br>Tsr1(3);<br><br>H6a(5);<br>H7(1);<br>H12(1);<br>H13(2);<br>H15(5); |
| H6 | 34 of 37<br>56-72;<br>76-92; | rpS6(17);<br>rpS24(3);<br><br>H8(5);<br>H10(4);<br>H13(3);<br>H15(6); | 34 of 37<br>56-72;<br>76-92; | rpS6(18);<br>rpS24(4);<br><br>H8(5);<br>H10(4);<br>H13(3);<br>H15(5); |

| | | | | |
|---|---|---|---|---|
| H6a | 11 of 11<br>93-99;<br>383-386; | rpS4(7);<br>rpS8(3);<br><br>H5(4);<br>H7(2);<br>H12(3);<br>H13(3);<br>H15(1); | 11 of 11<br>93-99;<br>383-386; | rpS4(7);<br>rpS8(3);<br><br>H5(4);<br>H7(2);<br>H12(3);<br>H13(3);<br>H15(1); |
| H7 | 50 of 58<br>100-129;<br>289-308; | rpS4(8);<br>rpS6(5);<br>rpS8(4);<br>rpS11(12);<br>Tsr1(1);<br><br>H5(1);<br>H6a(2);<br>H8(1);<br>H9_ES3a(2);<br>H9(6);<br>H11(6);<br>H12(3);<br>H13(1);<br>H21(2); | 50 of 58<br>100-129;<br>289-308; | rpS4(10);<br>rpS6(4);<br>rpS8(5);<br>rpS11(9);<br>Tsr1(1);<br><br>H5(1);<br>H6a(2);<br>H8(1);<br>H9_ES3a(2);<br>H9(6);<br>H11(6);<br>H12(2);<br>H13(1);<br>H21(2); |
| H8 | 38 of 38<br>138-175; | rpS6(31);<br>rpS24(6);<br><br>H6(5);<br>H7(1);<br>H9_ES3a(1);<br>H10(2);<br>H14(5);<br>H44(1); | 38 of 38<br>138-175; | rpS6(28);<br>rpS24(7);<br><br>H6(5);<br>H7(1);<br>H9_ES3a(1);<br>H10(2);<br>H14(6); |
| H9_ES3a | 19 of 28<br>176-187;<br>197-203; | rpS6(3);<br>rpS8(3);<br><br>H7(2);<br>H8(1);<br>H10(1); | 19 of 28<br>176-187;<br>197-203; | rpS6(3);<br>rpS8(2);<br><br>H7(2);<br>H8(1);<br>H10(1); |
| H9 | 38 of 38<br>204-221;<br>246-265; | rpS4(7);<br>rpS6(1);<br>rpS8(11);<br>rpS11(11);<br><br>H7(7);<br>H9_ES3b(3);<br>H10(1);<br>H21_ES6d(10); | 38 of 38<br>204-221;<br>246-265; | rpS4(8);<br>rpS6(1);<br>rpS8(11);<br>rpS11(11);<br><br>H7(7);<br>H9_ES3b(3);<br>H10(1);<br>H21_ES6d(10); |
| H9_ES3b | 4 of 23<br>241-244; | rpS4(1);<br>rpS6(2);<br><br>H9(4); | 5 of 23<br>222-222;<br>241-244; | rpS4(1);<br>rpS6(3);<br><br>H9(4);<br>H21_ES6d(3); |

| | | | | |
|---|---|---|---|---|
| H10 | 19 of 23<br>266-276;<br>281-288; | rpS6(5);<br><br>H6(2);<br>H8(6);<br>H9_ES3a(1)<br>H9(1); | 19 of 23<br>266-276;<br>281-288; | rpS6(5);<br><br>H6(2);<br>H8(6);<br>H9_ES3a(1);<br>H9(1); |
| H11 | 52 of 52<br>309-360; | rpS8(23);<br>rpS11(18);<br>rpS23(5);<br>Tsr1(3);<br><br>H7(6);<br>H12(1);<br>H20(1);<br>H27(3); | 52 of 52<br>309-360; | rpS8(22);<br>rpS11(15);<br>rpS23(4);<br>Tsr1(3);<br><br>H7(6);<br>H12(1);<br>H20(1);<br>H27(2); |
| H12 | 23 of 23<br>361-383; | rpS4(2);<br>rpS9(3);<br>rpS11(1);<br>rpS23(1);<br><br>H1(1);<br>H3(5);<br>H5(1);<br>H6a(2);<br>H7(2);<br>H11(1);<br>H19(3);<br>H21(4); | 23 of 23<br>361-383; | rpS4(2);<br>rpS9(3);<br>rpS11(1);<br>rpS23(1);<br><br>H1(1);<br>H3(3);<br>H5(1);<br>H6a(2);<br>H7(1);<br>H11(1);<br>H19(3);<br>H21(4); |
| H13 | 22 of 22<br>388-409; | rpS4(1);<br>rpS6(4);<br>rpS8(9);<br><br>H5(2);<br>H6(3);<br>H6a(1);<br>H7(1);<br>H14(2);<br>H44(7); | 22 of 22<br>388-409; | rpS4(1);<br>rpS6(3);<br>rpS8(9);<br><br>H5(1);<br>H6(3);<br>H6a(1);<br>H7(1);<br>H14(2);<br>H44(7); |
| H14 | 14 of 14<br>410-423; | rpS6(2);<br><br>H8(3);<br>H13(1); | 14 of 14<br>410-423; | rpS6(2);<br><br>H8(3);<br>H13(1); |
| H15 | 31 of 31<br>435-465; | rpS4(8);<br>rpS23(4);<br>rpS24(4);<br>Tsr1(1);<br><br>H3(1);<br>H4(2);<br>H5(13);<br>H6(9);<br>H6a(2);<br>H17(2); | 31 of 31<br>435-465; | rpS4(8);<br>rpS23(4);<br>rpS24(4);<br>Tsr1(1);<br><br>H3(1);<br>H4(2);<br>H5(13);<br>H6(9);<br>H6a(2);<br>H17(3); |
| H16 | 17 of 31<br>479-487;<br>501-505;<br>507-509; | rpS9(2);<br>rpS30(1);<br><br>H17(1); | 17 of 31<br>479-487;<br>501-505;<br>507-509; | rpS9(1);<br>rpS30(2);<br><br>H17(1); |

| | | | | |
|---|---|---|---|---|
| H17 | 34 of 34<br>510-543; | rpS9(10);<br>rpS24(11);<br>rpS30(2);<br>Tsr1(2);<br><br>H4(4);<br>H15(5);<br>H16(1);<br>H18(1); | 34 of 34<br>510-543; | rpS9(11);<br>rpS24(12);<br>Tsr1(1);<br><br>H4(3);<br>H15(6);<br>H16(1);<br>H18(1); |
| H18 | 51 of 51<br>544-594; | rpS9(4);<br>rpS23(10);<br>rpS30(7);<br><br>H1(4);<br>H3(3);<br>H4(6);<br>H17(2); | 47 of 51<br>544-564;<br>567-577;<br>580-594; | rpS9(4);<br>rpS23(10);<br>rpS30(9);<br><br>H1(2);<br>H3(3);<br>H4(5);<br>H17(2); |
| H19 | 20 of 20<br>607-622;<br>1104-1107; | rpS11(3);<br>rpS23(9);<br><br>H1(2);<br>H2(4);<br>H3(1);<br>H12(2);<br>H20(2);<br>H25(6);<br>H26a(4); | 20 of 20<br>607-622;<br>1104-1107; | rpS11(2);<br>rpS23(10);<br>Tsr1(3);<br><br>H1(3);<br>H2(4);<br>H12(2);<br>H20(3);<br>H25(6);<br>H26a(4); |
| H20 | 27 of 27<br>623-638;<br>966-976; | rpS7(3);<br>rpS11(1);<br>rpS13(6);<br>rpS22(6);<br>rpS23(3);<br><br>H11(1);<br>H22(5);<br>H23a(3);<br>H24(3);<br>H25(5); | 27 of 27<br>623-638;<br>966-976; | rpS7(3);<br>rpS11(1);<br>rpS13(5);<br>rpS22(6);<br>rpS23(2);<br><br>H11(1);<br>H22(5);<br>H23a(3);<br>H24(2);<br>H25(6); |
| H21_ES6c | 26 of 55<br>639-652;<br>682-693; | rpS7(7);<br>rpS22(2);<br><br>H21_ES6b(1); | 26 of 55<br>639-652;<br>682-693; | rpS7(8);<br>rpS22(2);<br><br>H21_ES6b(1); |
| H21_ES6b | 2 of 49<br>694-695; | rpS7(4);<br>1(2);<br>H21_ES6c(2); | 2 of 49<br>694-695; | rpS7(4);<br>1(2);<br>H21_ES6c(2); |
| H21 | 44 of 44<br>743-764;<br>789-810; | rpS4(8);<br>rpS7(3);<br>rpS9(4);<br>rpS22(9);<br><br>H7(2);<br>H12(4);<br>H21_unk(2);<br>H21_ES6a(1); | 44 of 44<br>743-764;<br>789-810; | rpS4(8);<br>rpS7(3);<br>rpS9(4);<br>rpS11(1);<br>rpS22(10);<br><br>H7(2);<br>H12(4);<br>H21_unk(2);<br>H21_ES6a(1); |

| | | | | |
|---|---|---|---|---|
| H21_unk | 9 of 9<br>765-773; | rpS4(3);<br>rpS9(14);<br>rpS24(1);<br><br>H4(2);<br>H21(2);<br>H21_ES6a(2); | 9 of 9<br>765-773; | rpS4(3);<br>rpS9(11);<br>rpS24(1);<br><br>H4(2);<br>H21(2);<br>H21_ES6a(2); |
| H21_ES6a | 12 of 15<br>774-779;<br>783-788; | rpS4(3);<br>rpS24(3);<br><br>H21(1);<br>H21_unk(2); | 12 of 15<br>774-779;<br>783-788; | rpS4(4);<br>rpS24(3);<br><br>H21(1);<br>H21_unk(2); |
| H21_ES6d | 44 of 49<br>811-833;<br>839-859; | rpS7(7);<br>rpS13(2);<br><br>H9(5); | 44 of 49<br>811-833;<br>839-859; | rpS7(7);<br>rpS13(2);<br><br>H9(5);<br>H9_ES3b(1); |
| H22 | 45 of 45<br>860-883;<br>945-965; | rpS1(3);<br>rpS7(1);<br>rpS13(34);<br>rpS22(4);<br>rpS27(9);<br><br>H20(4);<br>H23(2);<br>H23a(6);<br>H26(3); | 45 of 45<br>860-883;<br>945-965; | rpS1(3);<br>rpS7(1);<br>rpS13(32);<br>rpS22(4);<br>rpS27(10);<br><br>H20(4);<br>H23(2);<br>H23a(6);<br>H26(3); |
| H23 | 45 of 45<br>884-928; | rpS1(7);<br>rpS14(19);<br><br>H22(1);<br>H24(11); | 45 of 45<br>884-928; | rpS1(4);<br>rpS14(20);<br><br>H22(1);<br>H24(11); |
| H23a | 16 of 16<br>929-944; | rpS1(5);<br>rpS13(2);<br>rpS14(1);<br>Nob1(3);<br><br>H20(5);<br>H22(3);<br>H24(1);<br>H26(2); | 16 of 16<br>929-944; | rpS1(5);<br>rpS13(2);<br>rpS14(1);<br><br>H20(5);<br>H22(3);<br>H24(1);<br>H26(3); |
| H24 | 51 of 51<br>977-1027; | rpS13(1);<br>rpS14(3);<br>Pno1(5);<br><br>H20(2);<br>H23(10);<br>H23a(1);<br>H25(1);<br>H27(6);<br>H45(14); | 51 of 51<br>977-1027; | rpS13(1);<br>rpS14(3);<br>Pno1(5);<br>Tsr1(1);<br><br>H20(2);<br>H23(9);<br>H23a(1);<br>H25(1);<br>H27(5);<br>H45(13); |

| | | | | |
|---|---|---|---|---|
| H25 | 22 of 22<br>1028-1037;<br>1094-1105; | rpS2(3);<br>rpS13(2);<br>rpS22(11);<br>rpS23(7);<br><br>H19(7);<br>H20(5);<br>H24(1);<br>H26(1);<br>H26a(4);<br>H45(1); | 22 of 22<br>1028-1037;<br>1094-1105; | rpS13(2);<br>rpS22(11);<br>rpS23(7);<br><br>H19(7);<br>H20(6);<br>H24(1);<br>H26(1);<br>H26a(3);<br>H45(1); |
| H26 | 29 of 29<br>1038-1051;<br>1067-1081; | rpS0(1);<br>rpS1(5);<br>rpS13(1);<br>rpS22(1);<br>rpS27(4);<br>Nob1(12);<br><br>H22(5);<br>H23a(2);<br>H25(1);<br>H26_ES7(1);<br>H26a(3); | 29 of 29<br>1038-1051;<br>1067-1081; | rpS1(4);<br>rpS22(1);<br>rpS27(4);<br><br>H22(6);<br>H23a(2);<br>H25(1);<br>H26_ES7(1);<br>H26a(4); |
| H26_ES7 | 13 of 15<br>1052-1058;<br>1061-1066; | rpS1(8);<br><br>H26(2); | 13 of 15<br>1052-1058;<br>1061-1066; | rpS1(6);<br><br>H26(2); |
| H26a | 12 of 12<br>1082-1093; | rpS2(1);<br>rpS21(1);<br>Nob1(2);<br><br>H2(4);<br>H19(3);<br>H25(2);<br>H26(4);<br>H36(1); | 12 of 12<br>1082-1093; | <br>H2(4);<br>H19(3);<br>H25(2);<br>H26(4); |
| H27 | 28 of 28<br>1109-1136; | rpS23(3);<br>Tsr1(3);<br><br>H1(2);<br>H2(2);<br>H11(1);<br>H24(6);<br>H45(1); | 28 of 28<br>1109-1136; | rpS23(3);<br>Tsr1(6);<br><br>H1(2);<br>H2(2);<br>H11(1);<br>H24(6);<br>H45(1); |
| H28 | 30 of 38<br>1145-1148;<br>1152-1162;<br>1616-1626;<br>1629-1632; | rpS28(3);<br><br>H2(1);<br>H29(3);<br>H35(1);<br>H36(2);<br>H43(1); | 29 of 38<br>1145-1147;<br>1152-1162;<br>1616-1626;<br>1629-1632; | rpS28(4);<br>Pno1(4);<br><br>H2(2);<br>H29(2);<br>H43(1);<br>H44(1); |
| H29 | 14 of 14<br>1163-1169;<br>1575-1581; | rpS5(1);<br>Rio2(3);<br><br>H28(2);<br>H43(4); | 14 of 14<br>1163-1169;<br>1575-1581; | rpS5(2);<br>Rio2(4);<br><br>H28(1);<br>H43(4); |

| | | | | |
|---|---|---|---|---|
| H30 | 23 of 23<br>1170-1179;<br>1458-1470; | rpS15(2);<br>rpS19(1);<br>Rio2(2);<br><br>H31_unk1(2);<br>H31(1);<br>H32(1);<br>H42(8);<br>H43(2); | 23 of 23<br>1170-1179;<br>1458-1470; | rpS15(2);<br>rpS19(1);<br>Rio2(4);<br><br>H31_unk1(2);<br>H42(8);<br>H43(2); |
| H31_unk1 | 6 of 6<br>1180-1185; | rpS15(3);<br>Tsr1(3);<br><br>H30(2);<br>H32(5); | 6 of 6<br>1180-1185; | rpS15(5);<br>Tsr1(4);<br><br>H30(2);<br>H32(5); |
| H31 | 4 of 16<br>1186-1188;<br>1201-1201; | Tsr1(3);<br><br>H30(1);<br>H31_unk2(1);<br>H32(1);<br>H43(3); | 3 of 16<br>1186-1187;<br>1201-1201; | <br>H31_unk2(1);<br>H32(2);<br>H43(3); |
| H31_unk2 | 7 of 7<br>1202-1208; | H31(1);<br>H32(3);<br>H42(4);<br>H43(1); | 7 of 7<br>1202-1208; | H31(1);<br>H32(2);<br>H42(4);<br>H43(1); |
| H32 | 20 of 22<br>1209-1216;<br>1446-1457; | rpS15(5);<br>Enp1(2);<br>Tsr1(2);<br><br>H30(1);<br>H31_unk1(3);<br>H31(1);<br>H31_unk2(4);<br>H33(3);<br>H34(2); | 17 of 22<br>1209-1216;<br>1448-1456; | rpS15(4);<br>Enp1(1);<br>Tsr1(2);<br><br>H31_unk1(3);<br>H31(1);<br>H31_unk2(3);<br>H33(2); |
| H33 | 46 of 47<br>1219-1227;<br>1229-1265; | rpS15(4);<br><br>H32(6);<br>H34(4); | 42 of 47<br>1219-1227;<br>1229-1245;<br>1248-1256;<br>1258-1263;<br>1265-1265; | rpS15(4);<br>Enp1(1);<br><br>H32(3); |
| H34 | 20 of 46<br>1266-1271;<br>1278-1281;<br>1428-1430;<br>1439-1445; | Enp1(2);<br>Tsr1(1);<br><br>H32(2);<br>H33(3); | 0 of 46 | - |
| H35 | 15 of 15<br>1288-1293;<br>1321-1329; | rpS0(4);<br>rpS2(1);<br>Nob1(1);<br><br>H28(1);<br>H36(4);<br>H37(1); | 0 of 15 | - |

| | | | | |
|---|---|---|---|---|
| H36 | 12 of 12<br>1294-1305; | rpS0(3);<br>rpS2(2);<br>Nob1(2);<br><br>H2(4);<br>H26a(2);<br>H28(1);<br>H35(4); | 0 of 12 | - |
| H37 | 15 of 15<br>1306-1320; | rpS0(1);<br>Nob1(4);<br><br>H35(2); | 0 of 15 | - |
| H38 | 10 of 15<br>1332-1337;<br>1416-1419; | rpS16(1);<br><br>H40(2); | 0 of 15 | - |
| H39 | 29 of 33<br>1338-1339;<br>1341-1345;<br>1347-1353;<br>1372-1386; | rpS16(2);<br><br>H39_ES9(1);<br>H40(1);<br>H41(2); | 11 of 33<br>1347-1353;<br>1374-1377; | rpS16(1);<br><br>H41(2); |
| H39_ES9 | 16 of 16<br>1354-1369; | rpS16(3);<br>rpS19(8);<br><br>H39(1); | 14 of 16<br>1354-1361;<br>1364-1369; | rpS16(3);<br>rpS19(7); |
| H40 | 10 of 28<br>1387-1391;<br>1408-1412; | H38(1);<br>H39(1); | 0 of 28 | - |
| H41 | 66 of 66<br>1471-1536; | rpS5(12);<br>rpS16(6);<br>rpS18(3);<br>rpS19(26);<br>rpS25(7);<br><br>H39(2);<br>H42(7);<br>H43(8); | 66 of 66<br>1471-1536; | rpS5(12);<br>rpS16(6);<br>rpS18(3);<br>rpS19(29);<br>rpS25(7);<br><br>H39(1);<br>H42(5);<br>H43(9); |
| H42 | 38 of 38<br>1537-1574; | rpS5(1);<br>rpS15(14);<br>rpS18(14);<br>rpS19(3);<br>Rio2(4);<br><br>H30(6);<br>H31_unk2(2);<br>H41(5);<br>H43(3); | 38 of 38<br>1537-1574; | rpS5(1);<br>rpS15(15);<br>rpS18(16);<br>rpS19(2);<br>Rio2(4);<br><br>H30(6);<br>H31_unk2(2);<br>H41(5);<br>H43(3); |

| | | | | |
|---|---|---|---|---|
| H43 | 34 of 34<br>1582-1615; | rpS5(11);<br>rpS16(9);<br>rpS19(5);<br>rpS28(2);<br><br>H28(1);<br>H29(3);<br>H30(4);<br>H31(1);<br>H31_unk2(1);<br>H41(9);<br>H42(4); | 34 of 34<br>1582-1615; | rpS5(12);<br>rpS16(5);<br>rpS19(5);<br>rpS28(2);<br><br>H28(1);<br>H29(3);<br>H30(4);<br>H31(1);<br>H31_unk2(1);<br>H41(8);<br>H42(4); |
| H44 | 95 of 134<br>1646-1692;<br>1710-1754;<br>1767-1769; | rpS6(5);<br>rpS8(5);<br>rpS23(2);<br>Tsr1(7);<br><br>H8(1);<br>H13(7);<br>H45(1); | 94 of 134<br>1646-1692;<br>1710-1753;<br>1767-1769; | rpS6(4);<br>rpS8(7);<br>rpS23(2);<br>Tsr1(4);<br><br>H13(7);<br>H28(1); |
| H45 | 31 of 31<br>1770-1800; | rpS14(4);<br>Pno1(17);<br>Nob1(1);<br><br>H24(11);<br>H25(2);<br>H27(1);<br>H44(1); | 30 of 31<br>1770-1799; | rpS14(4);<br>Pno1(17);<br>Tsr1(8);<br><br>H24(11);<br>H25(2);<br>H27(1); |
