## Supplementary material for "Impact of the yeast S0/uS2-cluster ribosomal protein rpS21/eS21 on rRNA folding and the architecture of small ribosomal subunit precursors": S1 Raw Images

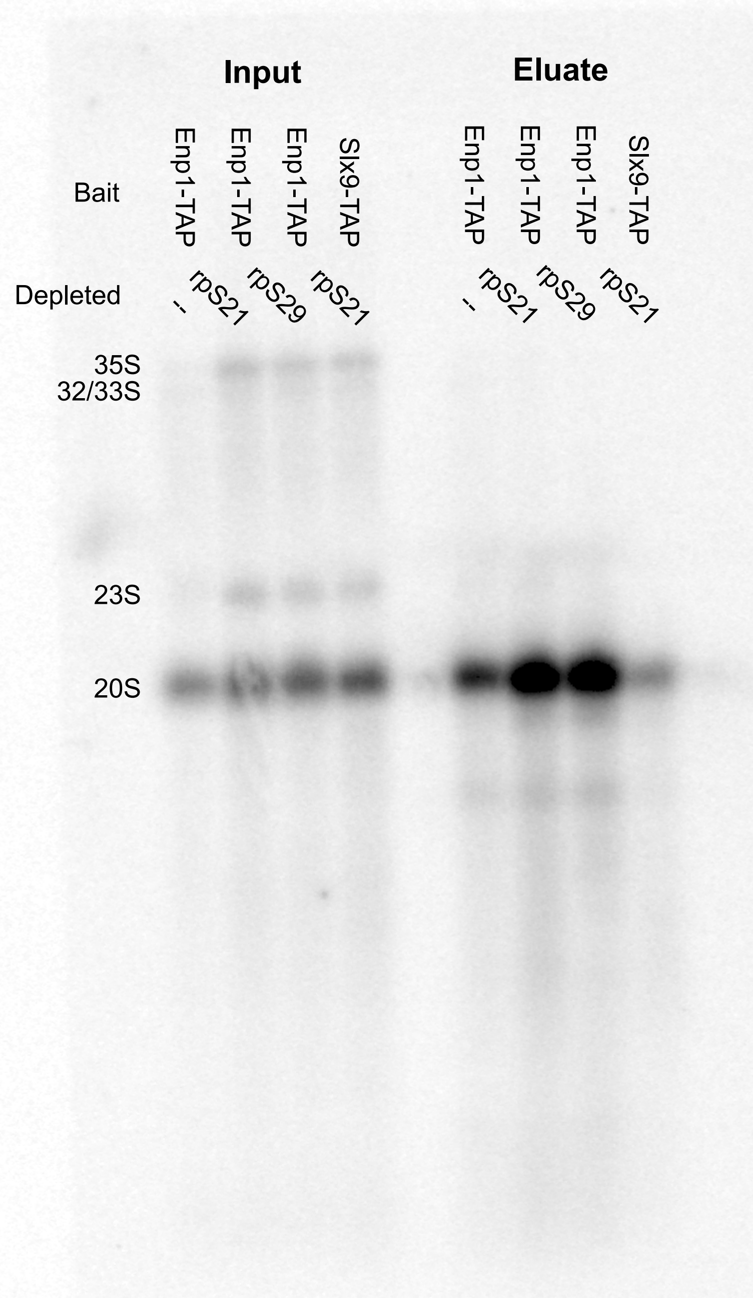


Blot exposition for S1 Figure A, probe O1819,

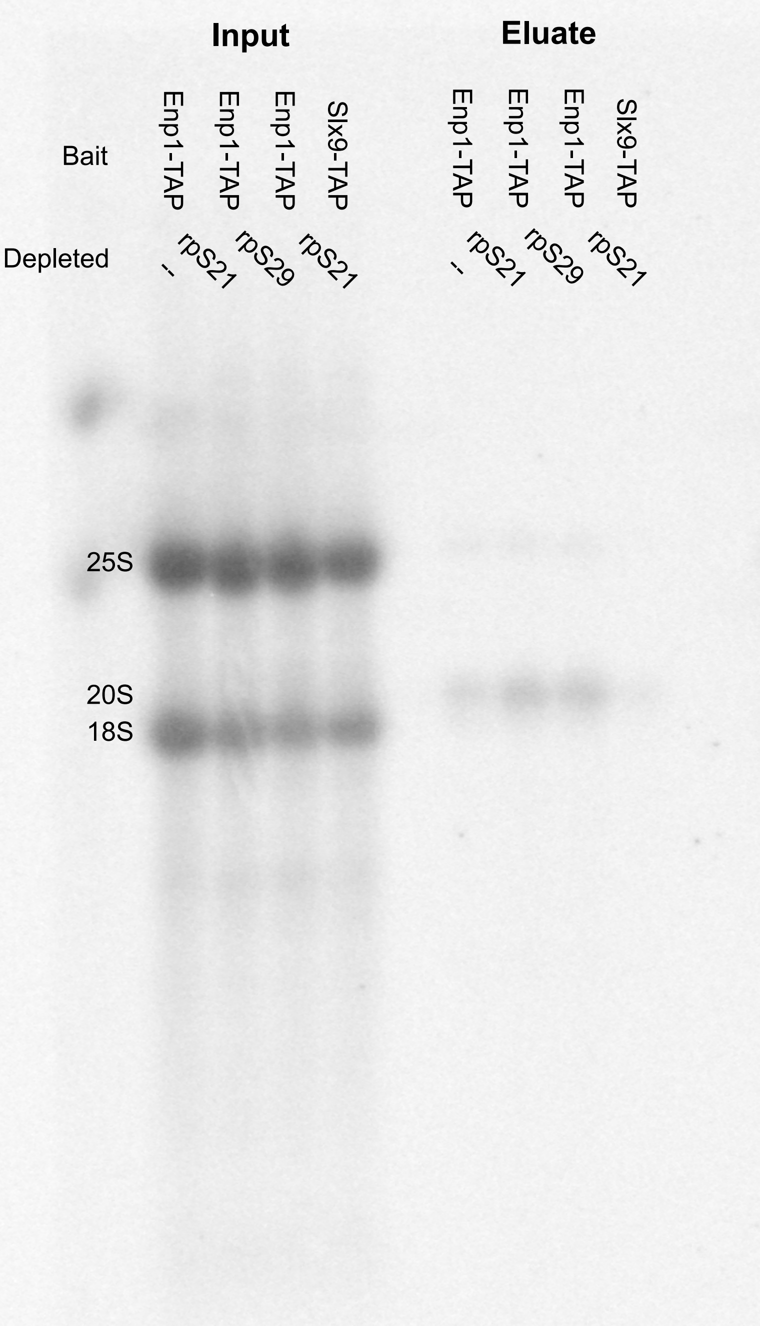


Blot exposition for S1 Figure B, probes O205+O212,

Images were generated by reading out Imaging plates exposed to 32P-labeled Northern blots with a Typhoon Imager FLA9500 (GE Healthcare)
